# A novel immunopeptidomic-based pipeline for the generation of personalized oncolytic cancer vaccines

**DOI:** 10.1101/2021.06.08.447483

**Authors:** Sara Feola, Jacopo Chiaro, Beatriz Martins, Salvatore Russo, Manlio Fusciello, Erkko Ylösmäki, Firas Hamdan, Michaela Feodoroff, Gabriella Antignani, Tapani Viitala, Sari Pesonen, Mikaela Grönholm, Rui M M Branca, Janne Lehtiö, Vincenzo Cerullo

## Abstract

Beside the isolation and identification of MHC-I restricted peptides from the surface of cancer cells, one of the challenges is eliciting an effective anti-tumor CD8+ T cell mediated response as part of therapeutic cancer vaccine. Therefore, the establishment of a solid pipeline for the downstream selection of clinically relevant peptides and the subsequent creation of therapeutic cancer vaccines are of utmost importance. Indeed, the use of peptides for eliciting specific anti-tumor adaptive immunity is hindered by two main limitations: the efficient selection of the most optimal candidate peptides and the use of a highly immunogenic platform to combine with the peptides to induce effective tumor-specific adaptive immune responses. Here, we describe for the first time a streamlined pipeline for the generation of personalized cancer vaccines starting from the isolation and selection of the most immunogenic peptide candidates expressed on the tumor cells and ending in the generation of efficient therapeutic oncolytic cancer vaccines. This immunopeptidomics-based pipeline was carefully validated in a murine colon tumor model CT26. Specifically, we used state-of-the-art immunoprecipitation and mass spectrometric methodologies to isolate >8000 peptide targets from the CT26 tumor cell line. The selection of the target candidates was then based on two separate approaches: RNAseq analysis and the HEX software. The latter is a tool previously developed by Chiaro et al. (1), able to identify tumor antigens similar to pathogen antigens, in order to exploit molecular mimicry and tumor pathogen cross-reactive T-cells in cancer vaccine development. The generated list of candidates (twenty-six in total) was further tested in a functional characterization assay using interferon-γ ELISpot (Enzyme-Linked Immunospot), reducing the number of candidates to six. These peptides were then tested in our previously described oncolytic cancer vaccine platform PeptiCRAd, a vaccine platform that combines an immunogenic oncolytic adenovirus (OAd) coated with tumor antigen peptides. In our work, PeptiCRAd was successfully used for the treatment of mice bearing CT26, controlling the primary malignant lesion and most importantly a secondary, non-treated, cancer lesion. These results confirmed the feasibility of applying the described pipeline for the selection of peptide candidates and generation of therapeutic oncolytic cancer vaccine, filling a gap in the field of cancer immunotherapy, and paving the way to translate our pipeline into human therapeutic approach.

## INTRODUCTION

The ligandome describes the peptide composition bound to the major histocompatibility complex (MHC) I and II presented on the cellular surface (2). Once being identified as targets by the immune system, the peptides in the MHC-I are the contact point between cytotoxic CD8+ T cells and the tumor cells. Thus, the knowledge of those peptides is a key point in designing therapeutic cancer vaccines to generate and stimulate specific anti-tumor adaptive immune responses. Moreover, the interest in identifying and exploiting these targets gained momentum following the breakthrough of the immune checkpoint inhibitors (ICIs), as it became clear that ICI treatment can unleash the specific anti-tumor T cell responses against these immunogenic candidate targets.(3). Indeed, the ICI therapy activates a pre-existing antitumor immune response with immune cell infiltration in the cancer lesions, defined as “hot” tumors; instead, tumors not infiltrated with immune cells are called “cold”. As a result, the response rate to the ICI therapy can vary from 40%-70% to 10-25% either due to the lack of immune cell infiltration into the tumor or other immunosuppressive mechanisms in the tumor microenvironment (TME) (4, 5). Currently, there is an urgent need to find a way to turn “cold” tumors to “hot” ones, making the ICI therapies more effective. In this context, the development of effective peptide-based cancer vaccines for therapeutic approaches is facing two main challenges: the criteria to select peptides able to elicit an immune response and the use of an adjuvant to increase the anti-tumor immune response of the immunizing peptides. In the present work, to overcome these issues, we have developed a pipeline that covers the diverse developmental stages of therapeutic cancer vaccines, moving from the isolation of the MHC-I restricted tumor peptides, to the selection and screening of target candidates until the generation of an oncolytic cancer vaccine. First, we selected the known murine immunogenic tumor model CT26, allowing the study of the anti-tumor response (6). We investigated the MHC-I antigen landscape of CT26 applying state-of-art immunopeptidome and mass spectrometric methodologies. The immunopeptidome profile was carefully analyzed and found to be qualitatively in line with already published dataset; the result list of peptides was then investigated through two approaches: RNA seq and the HEX software. The latter is a tool that identifies tumor antigens similar to pathogen antigens, exploiting the cross-mimicry and cross-reactive T cells for clinical applications (1). The peptides derived from those analyses were then investigated *in vivo*, by pre-immunizing mice with the adjuvant poly:(IC) and the peptides; the splenocytes were then harvested and functional characterization was performed by interferon-γ ELISpot (Enzyme-Linked Immunospot), deconvoluting the single peptide immunogenicity. For the last part of our pipeline, after the functional characterization, the selected peptides were used to generate an oncolytic cancer vaccine. To take full advantage of viral immunogenicity to induce a specific anti-tumor T cell response, we used our previously developed platform, PeptiCRAd based on an OAd coated with immunogenic tumor antigen peptides (7, 8). The peptide candidates in this study were tested in our PeptiCRAd platform, which in the present work consisted of a conditionally replicating OAd armed with two immune activating ligands, the ligand for cluster of differentiation 40 (CD40L) and the ligand for tumor necrosis factor receptor superfamily member 4 (OX40L), named VALO-mD901 (9). Intratumoral administration of PeptiCRAd coated with the peptides selected based on our pipeline, controlled the tumor growth in CT26 tumor bearing mice. Additionally, we observed that the specific anti-tumor immune activation generated in the primary tumor could be extended to a second tumor lesion, in a phenomenon known as “abscopal effect”. Thus, we developed and validated a pipeline moving from the isolation of the peptides to the selection of the target candidates until the combination of these in our PeptiCRAd platform, showing the efficacy in a pre-clinical model of colon cancer on to the primary tumor and distant lesions. To the best of our knowledge, the described pipeline covers for the first time all the stages of a personalized therapeutic cancer vaccine development, starting from the isolation of MHC-I restricted peptides derived from the primary tumor to their analysis *in silico* and *in vivo* to identify the best target candidates. Finally, an OAd was coated with these peptides to generate an effective therapeutic cancer vaccine. The pipeline can be translated to personalized cancer treatment in relevant clinical application as the OAd can be easily coated with the unique repertoire of patient-specific tumor peptides profile, a prerequisite for personalized therapy.

## MATERIALS AND METHODS

### Cell lines and reagents

Murine colon carcinoma CT26 cell line was purchased from ATCC (ATTC CRL-2639) and cultured in RPMI-1640 supplemented with 1% GlutaMAX (GIBCO, Invitrogen, Carlsbad, CA, USA), 10% heat inactivated fetal bovine serum (HI-FBS, GIBCO, Invitrogen, Carlsbad, CA, USA) and 1% Penicillin-Streptomycin (10,000 U/mL) (GIBCO, Invitrogen, Carlsbad, CA, USA). The cells were cultivated in 37°C, 5% CO_2_ in a humidified atmosphere.

Poly(I:C) (HMW) VacciGrade 10mg was obtained from Invivogen (San Diego, California). The following peptides were used for the pre-immunization experiment:

SYHPALNAI, SYLTSASSL, YYVRILSTI, SYLPPGTSL, RYLPAPTAL, KYIPAARHL, AFHSSRTSL, NYNSVNTRM, SYSDMKRAL, FYEKNKTLV, KGPNRGVII, FYKNGRLAV, LYKESLSRL, SYRDVIQEL, KFYDSKETV, KYLNVREAV, HYLPDLHHM, SGPNRFILI, SYIIGTSSV, RGPYVYREF, FYATIIHDL, GYMTPGLTV, SYLIGRQKI, AGASRIIGI, QGPEYIERL, SYIHQRYIL.

All peptides were purchased from Zhejiang Ontores Biotechnologies (Zhejiang, China).

The following peptides were used through the animal study and purchased from PepScan (LelyStand, the Netherlands): KKKKKKSYLPPGTSL (Mavs), KKKKKKRYLPAPTAL (Fanca), KKKKKKYIPAARHL (Zw10), KKKKKKLYKESLSRL (Myh14), KKKKKKYLNVREAV (Chac1), KKKKKKKFYATIIHDL (Ndst3), SYLPPGTSL (Mavs), RYLPAPTAL (Fanca), KYIPAARHL (Zw10), LYKESLSRL (Myh14), KYLNVREAV (Chac1), FYATIIHDL (Ndst3).

### Oncolytic Adenovirus

In this study the virus VALO-mD901 was used, and it was generated according to Ylösmäki et al. (9). Briefly, VALO-mD901 is a conditionally replicating adenovirus serotype 5 with adenovirus 3 fiber knob modification and 24-base pair deletion of the gene E1A. The E3 region was replaced with human cytomegalovirus (CMV) promoter region, murine OX40L, 2A self-cleaving peptide sequence, murine CD40Lgene and β-rabbit globin polyadenylation signal. The viral particle (VP) concentration was measured at 260nm, and infections units (IU) were determined by immunocytochemistry (ICC) by staining the hexon protein on infected A549 cells.

### IFN-γ ELISpot

IFN-γ ELISpot assays were performed using a commercially available mouse ELISpot reagent set (ImmunoSpot, Bonn Germany) and 20 ng/uL of each peptide was tested in *in vitro* stimulations of 3×10^5^ splenocytes for each well at 37 °C for 72h. Spots were counted using an ELISpot reader system (ImmunoSpot, Bonn Germany).

### PeptiCRAd complex formation

The PeptiCRAd complex was prepared by mixing the oncolytic adenovirus VALO-mD901 and each peptide with a polyK tail. We mixed polyK-extended epitopes with VALO-mD901 for 15 minutes at room temperature prior to treatments with the PeptiCRAd complexes. More details about the stability and formation of the complex can be found in our previous study (7).

### Animal experiment

All animal experiments were reviewed and approved by the Experimental Animal Committee of the University of Helsinki and the Provincial Government of Southern Finland (license number ESAVI/11895/2019).4-6 weeks old female Balb/cOlaHsd mice were obtained from Envigo (Laboratory, Bar Harbor, Maine UK).

For the pre-immunization experiment, mice (n=3 per group) were allocated in 9 different groups and each mouse was injected three times (one injection for each peptide) in three different areas (each injection contained 25ug of peptide+25ug of Poly I:C). The prime and boosting were done respectively at day 0 and 7 and the mice were sacrificed at day 14 For the tumor bearing mice experiment, 1×10^6^ and 6×10^5^ CT26 cells were injected subcutaneously into the right and in the left flank respectively. Details about the schedule of the treatment can be found in the figure legends. Viral dose was 1×10^9^ vp/tumor complexed with 20 µg of a single peptide or with 10 µg+10 µg mixture of two peptides.

### Flow Cytometry

The antibodies were: TruStain FcX™ anti-mouse CD16/32 (BioLegend), APC-H2Kd (BioLegend), BV711-CD3 (BD Horizon), PE-CF594-CD4 (BD Horizon), FITC-NK1.1 (Invitrogen), PE-PD1(BioLegend), APC-CXCR3 (BD Pharmigen), PE-CY7-TIM3 (BioLegend), BV510-CD8 (BD Horizon), V450-CXCR4 (BD Horizon). The data were acquired using BDLSRFORTESSA flow cytometer and analysed using FlowJo software v9 (Ashland, Oregon, USA).

### Purification and concentration of MHC-I peptides

MHC class I peptides were immunoaffinity purified from the CT26 mouse cell line using anti-mouse MHC class I (clone 34-1-2S, BioXcell, BE0180 Lebanon, USA). For sample preparation, the snap-frozen cell pellet (1×10^8^ cells for each replicate, in total 6 replicates) was incubated for 2h at 4°C in lysis buffer. The lysis buffer contained 150 mM NaCl, 50 mM TRIS-HCl, pH 7.4, protease inhibitors (A32955 Thermo Scientific Pierce, Waltham, Massachusetts, USA) and 1% Igepal (I8896 Sigma Aldrich, St. Louis, Missouri, USA). The lysates were first cleared by low-speed centrifugation for 10 min at 500xg, and then the supernatant was centrifuged for 30 min at 25,000xg. Next, MHC-I complexes were immunoaffinity purified loading the cleared lysate to the immunoaffinity column (AminoLink Plus Immobilization, Pierce) with covalently linked antibody according to the manufactureŕs instructions. Following binding, the affinity column was washed using 7 column volumes of each buffer (150 mM NaCl, 20 mM Tris-HCl; 400 mM NaCl, 20 mM Tris-HCl; 150 mM NaCl, 20 mM Tris-HCl and 20 mM TrisHCl, pH 8.0) and bound complexes were eluted in 0.1N acetic acid. Eluted HLA peptides and the subunits of the HLA complexes were desalted using SepPac-C18 cartridges (Waters) according to the protocol previously described by Bassani et al.(10). Briefly, the cartridge was prewashed with 80% acetonitrile in 0.1% trifluoroacetic acid (TFA) and then with 0.1% TFA. The peptides were purified from the MHC-I complex by elution with 30% acetonitrile in 0.1% TFA. Finally, the samples were dried using vacuum centrifugation (Eppendorf).

### Algorithms used for prediction of peptide ligands

Affinity to the H2Kd/H2Dd alleles was predicted for all eluted peptides identified in the CT26 cell line using NetMHC4.0 (11, 12). The threshold for binding was set to rank 2% to include only the binding partners.

### GIBBS clustering analysis

Clustering of peptides into groups based on sequence similarities was performed using the GibbsCluster-2.0 tool with the default settings (13, 14).

### Gene Ontology (GO) Enrichment Analysis

ClusterProfiler Bioconductor package (v. 3.12.0) in the RStudio server environment (v. 3.6.0) (15) was used for the functional annotation and visualization. ClusterProfiler implements a hypergeometric test to evaluate the statistical enrichment of the input gene list over the desired functional classes.

### Differential gene expression (DESeq) profile

Raw sequence data for colon tissue (source: GEO accession #GSE92563) and mTEC/CT26 (source: GEO accession: #GSE111092) were mapped to the mouse genome Mus_musculus GRCm38.95 using the online tool Chipster (16).

Briefly, fastaq files were combined for each sample sequencing using the function “Make a list of file names: paired end data”. The alignment to the reference genome and the count aligned reads per gene was done respectively with HISAT2 and HTSeq. Finally, the differential expression analysis used DESeq2, applying a cutoff for the adjusted p-value of 0.05 (Benjamini-Hochberg adjusted p-value). The “MultiQC function” was used to assess the quality of the fastaq files.

### LC-MS analysis of MHC-I peptides

Each dry sample was dissolved in 10 μL of LCMS solvent A (0.1% formic acid) by dispensing/aspirating 20 times with the micropipette. The nanoElute LC system (Bruker, Bremen, Germany) injected and loaded the 10 μl of sample directly onto the analytical column (Aurora C18, 25 cm long, 75 µm ID, 1.6 µm bead size, Ionopticks, Melbourne, Australia) constantly kept at 50℃ by a heating oven (PRSO-V2 oven, Sonation, Biberach, Germany). After washing and loading sample at a constant pressure of 800 bar, the LC system started a 30 min gradient from 0 to 32% solvent B (acetonitrile, 0.1% formic acid), followed by increase to 95% B in 5 min, and finally a wash of 10 min at 95% B, all at a flow rate of 300 nL/min. Online LC-MS was performed using a Tims TOF Pro mass spectrometer (Bruker, Bremen, Germany) with the CaptiveSpray source, capillary voltage 1500V, dry gas flow of 3L/min, dry gas temperature at 180℃. MS data reduction was enabled. Mass Spectra peak detection maximum intensity was set to 10. Mobilogram peak detection intensity threshold was set to 5000. Mass range was 300-1100 m/z, and mobility range was 0.6-1.30 V.s/cm^2^. MS/MS was used with 3 PASEF (Parallel Accumulation – Serial Fragmentation) scans (300ms each) per cycle with a target intensity of 20000 and intensity threshold of 1000, considering charge states 0-5. Active exclusion was used with release after 0.4 min, reconsidering precursor if current intensity is >4 fold the previous intensity, and a mass width of 0.015 m/z and a 1/k0 width of 0.015 V.s/cm^2^. Isolation width was defined as 2.00 m/z for mass 700 m/z and 3.00 m/z for mass 800 m/z. Collision energy was set as 10.62 eV for 1/k0 0.60 V.s/cm2 and 51.46 eV for 1/k0 1.30 V.s/cm^2^. Precursor ions were selected using 1 MS repetition and a cycle overlap of 1 with the default intensities/repetitions schedule.

### Proteomics database search

All MS/MS spectra were searched by PEAKS Studio X+ (v10.5 build 20191016) using a target-decoy strategy. The database used was the Swissprot Mouse protein database (including isoforms, 25284 entries, downloaded from uniprot.org on 20191127).

A precursor mass tolerance of 20 ppm and a product mass tolerance of 0.02 Da for CID-ITMS2 were used. Enzyme was none, digest mode unspecific, and oxidation of methionine was used as variable modification, with max 3 oxidations per peptide. A false discovery rate (FDR) cut-off of 1% was employed at the peptide level. The mass spectrometry proteomics data have been deposited to the ProteomeXchange Consortium via the PRIDE partner repository with the dataset identifier PXD026463. The dataset is currently hidden but will be made public upon eventual acceptance of the current manuscript.

### Surface Plasmon Resonance

Measurements were performed using a multi-parametric SPR Navi™ 220A instrument (Bionavis Ltd, Tampere, Finland). Phosphate buffered saline (PBS) (pH 7.4) was used as a running buffer, a constant flow rate of 20 µL/min was used throughout the experiments, and temperature was set to +20°C. Laser light with a wavelength of 670 nm was used for surface plasmon excitation and analysis. APTES ((3-aminopropyl) triethoxysilane) coated Au-SiO_2_ sensor slides were used to immobilize VALO-mD901 viruses on the sensors for evaluating peptide affinity and for assessing the number of peptides per VALO-mD901 virus. The APTES coated Au-SiO_2_ was prepared by first activating its surface by 5 min of oxygen plasma treatment followed by incubating the sensor in 50 mM APTES in isopropanol for 4 h, thus rendering the SPR sensor highly positively charged. The sensor was then washed and placed into the SPR device. The VALO-mD901 viruses were immobilized *in situ* on the sensor surface by injecting approximately 4.96×10^^11^vp/ml in PBS (pH 7.4) for 10 min, followed by a 10-min wash with PBS. For testing the interaction between various peptides and the immobilized VALO-mD901 viruses, 100 µM of the tested peptides were injected onto the viruses.

The SPR responses measured during virus immobilization as well as peptide interactions was used to estimate how many peptides were adsorbed per virus. This estimation is based on geometrical calculations including the SPR detection area (A_S_ = πr^2^, where r = 0.5 mm), diameter of the virus (d = 100 nm), the footprint area one virus covers on the SPR sensor (A_V_ = πr^2^, where r = 50 nm), the SPR signal response for a sensor fully covered with viruses (Δ° = 1.4°), the per cent coverage of viruses in the detection area (C(%) = (Measured SPR response)/(SPR response for full layer of viruses, i.e. 1.4°)), area covered by viruses in the detection area (A_V,cov_ = A_S_ × C(%)), number of viruses in detection area (N_V_ = A_V, cov_ / A_V_), mass/area of peptides determined from the corresponding SPR response (m/A = (Measured SPR response × 660 ng/cm^2^), mass of peptides in the detection area (m_P_ = m/A × A_S_) and the number of peptides in the detection area (N_P_ = [(m_P_/M_P_) × N_A_], where M_P_ is the molecular weight of the peptide and N_A_ is the Avogadro constant).

### Statistical Analysis

Statistical analysis was performed using GraphPad Prism 9.0 software (GraphPad Software Inc.). Details about the statistical tests for each experiment can be found in the corresponding figure legends.

## RESULTS

### Immunopeptidomic analysis reveals the MHC-I profile in a pre-clinical model of colon cancer

The identification and selection of candidate targets followed by the generation of therapeutic cancer vaccines is a scattered rather than a complete workflow. This drawback prompted us to develop a comprehensive pipeline that could cover the major steps in the process. First, we aimed to directly isolate MHC-I restricted peptides from the tumor surface as they are the key contact points between the tumor cells and the cytotoxic CD8+ T cells (Fig.1 **Step1)**. Next, the peptides were analyzed by mass-spectrometry (Fig.1 **Step2)** and the generated list of peptides was investigated with two independent approaches: RNAseq analysis and HEX software (Fig.1 **Step 3**). The selected peptides were then functionally characterized for their immunogenicity profile *in vivo* by ELIspot (Fig.1 **Step 4)** and the best candidates were modified to contain polyK attachment moiety and were analyzed by Surface Plasmon Resonance (SPR) for their binding affinity to the OAd (Fig.1 **Step 5)**. Finally, the peptides were used in our PeptiCRAd cancer vaccine platform (Fig.1 **Step 6**). As we sought to investigate whether the proposed pipeline could be applied for the development of therapeutic cancer vaccines, we selected the known immunogenic model CT26 (6), that expresses high surface level of MHC-I as shown in our flow-cytometry data (Suppl. Figure 1). We immunopurified MHC-I restricted peptides and analyzed the eluted peptides by tandem mass spectrometry. By using the murine reference proteome and applying an FDR threshold of 5% for peptide identification, a total of 8834 unique peptides were identified (Fig 2A). In order to assess the overall performance of the immunopurification of the MHC-I restricted peptides, we carefully investigated the presence of contaminants in the immunopurified peptides. Among those, the 7-13mers accounted for 5434 peptides (65% of the total eluted peptides) derived from 2218 unique source proteins (Fig. 2A). The peptides showed the typical aminoacid length distribution profile with the 9mers as the most enriched fraction, representing 21% of the total amount of peptides (Fig.2B). Next, the analysis of binding affinity to MHC-I showed that 81% (1413 of 1752) of 9mers were binders either for H2K^d^ or H2D^d^ (according to NetMHC4.0, applied rank <2%) with 62% of the binders showing preference for the H2K^d^ allele (Fig. 2C). Moreover, Gibbs analysis was used to deconvolute the consensus binding motifs of respective MHC-I alleles from the eluted 9mer peptides; these clustered in two distinct groups, with a preference for reduced amino acid complexity for residues at positions P2 and P9, matching remarkably well with the known motifs for H2K^d^ and H2D^d^ (Fig.2D). Overall, the analysis outcome was similar to published dataset (17) (aminoacidic length distribution, Gibbs clustering profile, amount of binders) confirming the good quality of the ligandome landscape identified.

**Figure 1.**
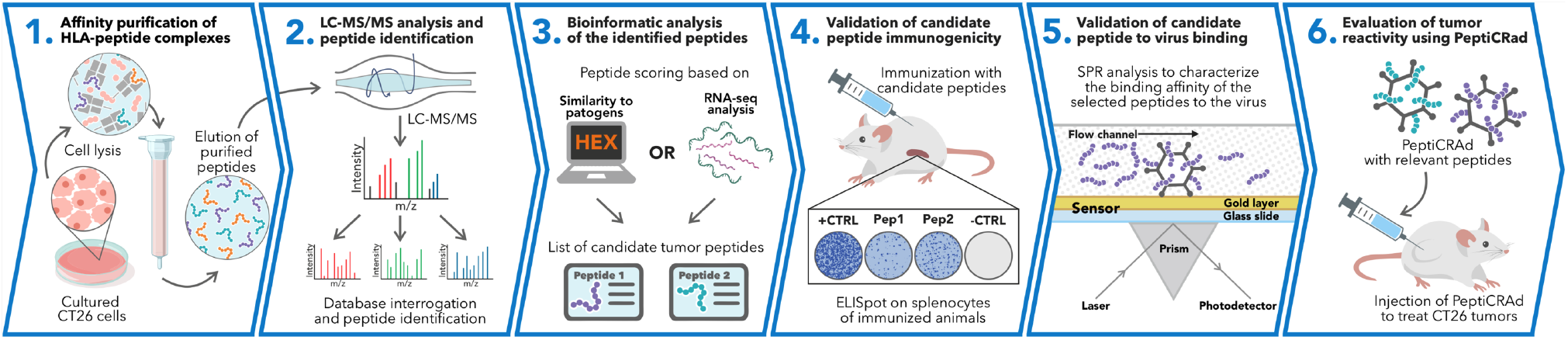
Schematic of the proposed immunopetidomic based pipeline. MHC-I peptides are immunopurified from the surface of tumor cells (**Step1**). Next, the peptides are analyzed by mass-spectrometry (**Step2**) and the generated list of peptides is investigated with two main approaches: RNAseq analysis and HEX software (**Step 3**). The selected peptides are then going through a functional characterization for their immunogenicity profile *in vivo* through ELISPOT assay (**Step 4**) and the best candidates are poly-lysine modified and analyzed by Surface Plasmon Resonance (SPR) for their binding affinity to the oncolytic adenovirus, OAd (**Step 5**). Finally, the peptides are used to decorate OAd to generate therapeutic cancer vaccine (PeptiCRAd) and tested in tumor bearing mice (**Step 6**).

**Figure 2.**
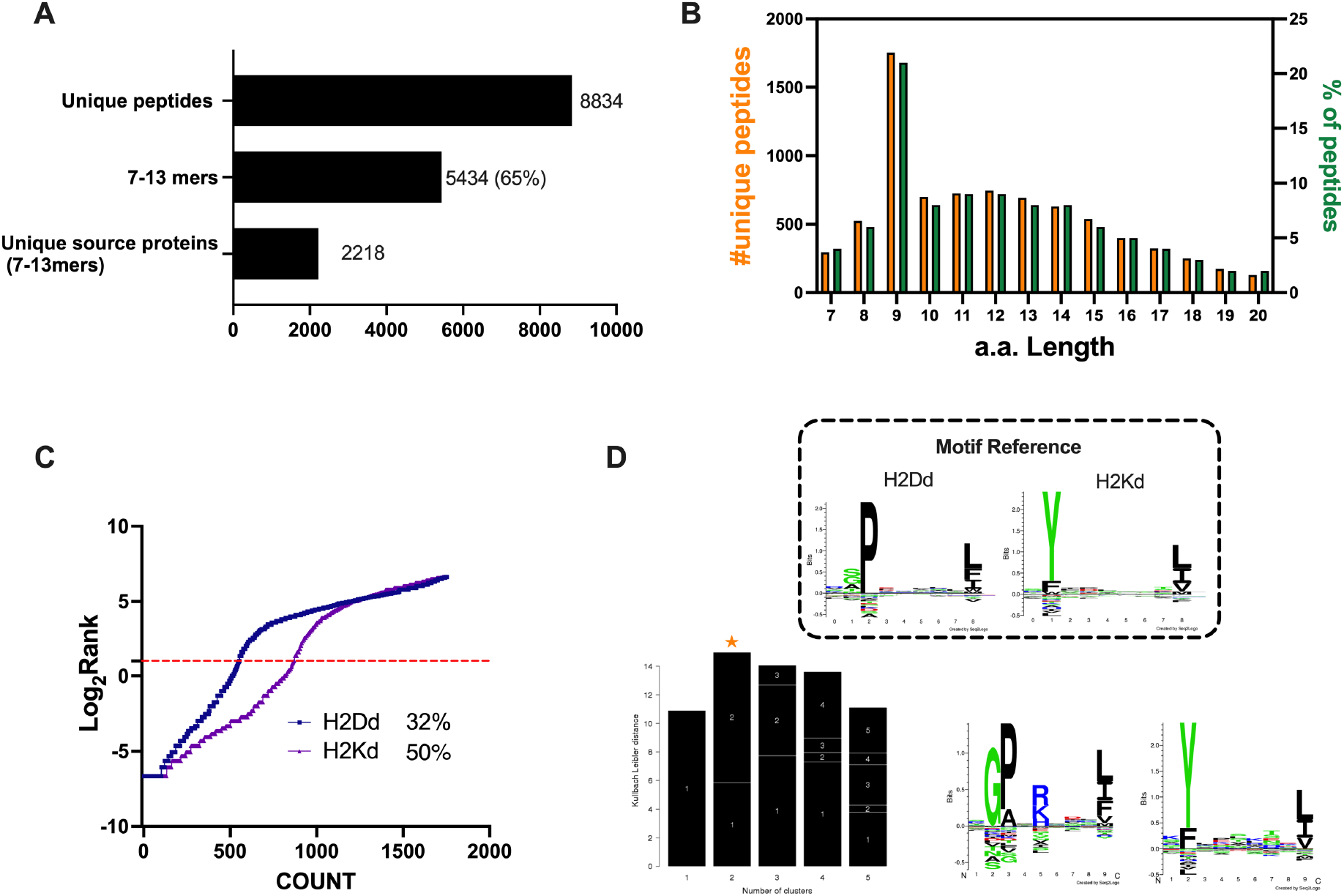
Properties of the peptides eluted from the CT26 tumor model. **A)** Unique peptides, 7-13 specimen and their respective source proteins are reported as finite number and depicted as bar plots. **B)** Overall peptides aminoacid length distribution is shown as function of number (left y axis) and percentage of occurrence (right y axis) **C)** The eluted 9mers were analyzed in regard to their binding affinity to H2K^d^ and H2D^d^. Binders and not binders were defined in NetMHCpan 4.0 Server (applied rank 2%). **D)** MHC-I consensus binding motifs. The consensus binding motifs among the eluted 9mers peptides was deconvoluted through Gibbs clustering analysis. The reference motif (according to NetMHCpan motif viewer) is depicted in the upper square. The clusters with the optimal fitness (higher KLD values, orange star) are shown and the sequence logo is represented.

Then, we aimed to investigate whether the MHC-I source proteins identified among the binders (9mers) were attributable to a specific biological process. Indeed, MHC-I peptides predominantly derived from cytosolic/nuclear proteins, that normally do not intersect the endocytic compartment and are mainly involved in maintaining the structure of the cell (cell proliferation, differentiation, signaling, translation) (18). To this end we performed a gene ontology enrichment analysis. As expected, the biological process highlighted the enrichment in pathways that comprise regulation of chromosome organization, DNA repair, ribosome biogenesis, RNA splicing, DNA-protein interactions and cytoskeleton organization (Fig.3A). Moreover, the linkage between the genes and the biological process depicted an overrepresentation of epigenetic regulators (e.g., histones, DNMT1) (Fig.3B **and** Suppl. Fig. 2), in line with preceding reports in literature (19). The cellular component (CC) and the molecular functions (MF) confirmed the nature of the source proteins, showing an enrichment for instance in nucleosome and chaperone proteins respectively; these are well known sources of MHC-I ligands (Suppl.Fig.3).

**Figure 3.**
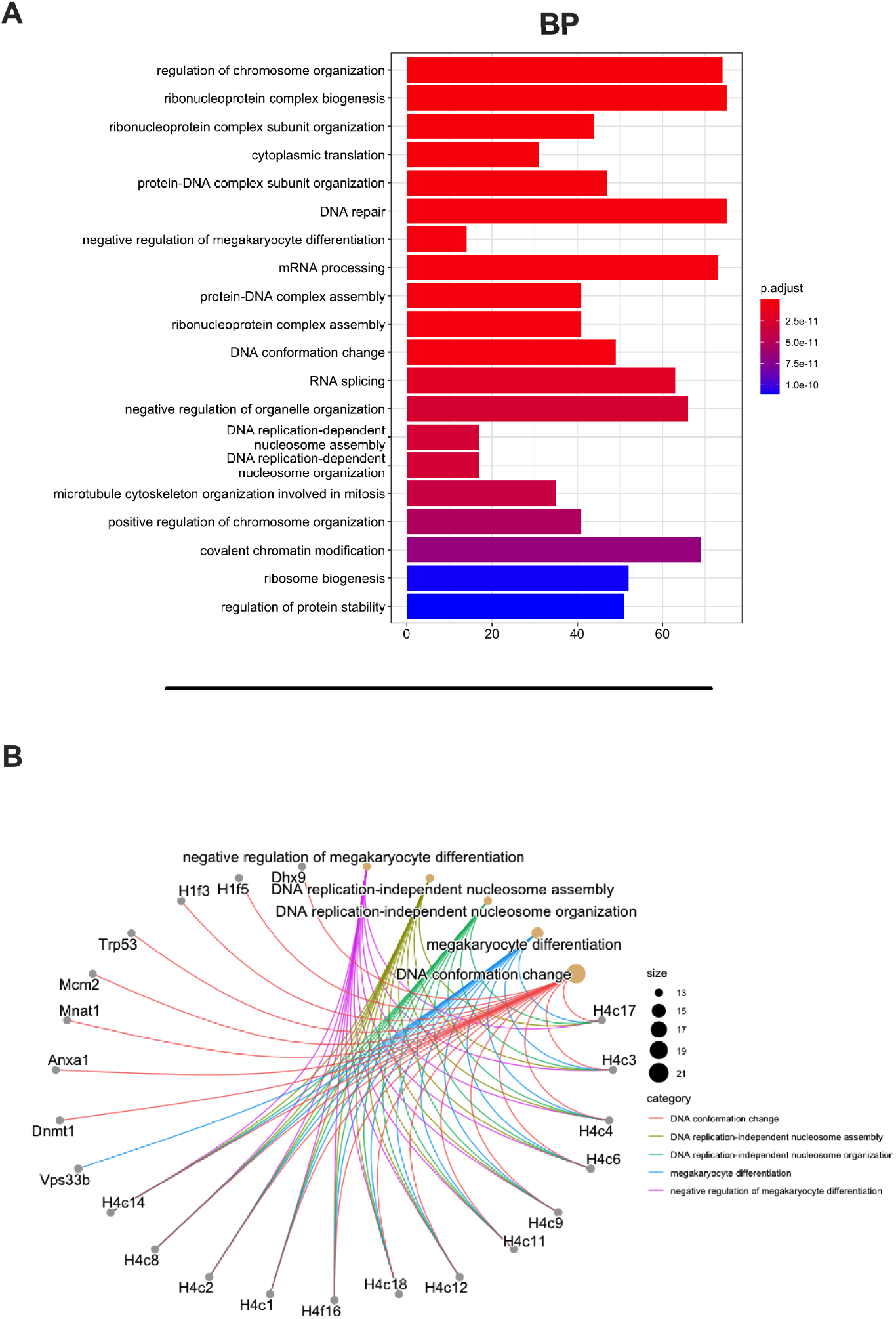
Gene Ontology (GO) enrichment analysis of the source proteins. **A)** GO enrichment was evaluated at Biological Process (BP); adjusted p-values of the first 20 statically relevant BP are depicted as color gradient and the respective number of genes is shown as bar plots. **B)** Genes and biological process linkages are summarized in a cnetplot graph. Each color line represents a different biological process category, and the bubble size symbolizes the number of genes.

Overall, these analyses assessed and demonstrated the reliability of the generated ligandome data set, confirming the robustness of the peptides list as true ligands and allowing us to proceed further with the downstream applications.

### *In silico* prediction of candidate targets based on RNAseq analysis and similarity to pathogen antigens

We carefully examined the list of generated peptides to check for the presence of contaminants and based on the aforementioned analysis, the eluted peptides resembled the MHC-I ligandome landscape. As we sought to generate and develop an effective therapeutic cancer vaccine, we next moved to selecting the best peptide candidates that could elicit a strong adaptive immune response. However, the criteria for selecting and narrowing down the number of peptide targets is still challenging for the field, usually involving laborious and time-consuming approaches and remaining therefore a critical question to address (20). To overcome this limitation, we analyzed the list of peptides adopting two parallel approaches. The first one is based on the RNA expression level of the source proteins of the MHC-I ligands. With this mind, we first identified the transcripts (and thus the corresponding source proteins) overrepresented in CT26 tumor cell line compared to normal cells. The RNA seq profile of the syngeneic mTEC (medullary thymic epithelial cells) and the colon Balb/c was used as normal control. Thus, we analyzed the differential gene expression (DESeq) profile between the CT26 and mTEC (Fig.4A) and CT26 and colon (Fig.4B) (standard cut-off values of fold-change 1.5 and a padj-value of 0.05, red square); then, we searched the source proteins of the 9mers ligands derived from our previously generated ligandome data set **(red dots in** Fig.4A-B) in the DESeq data for each expression profile analysis. In order to identify tumor associated antigens (TAA), we selected the liagandome source proteins for which the corresponding transcripts were overexpressed in both DESeq analyses (Fig.4A**-B, red dots within the red square**). Finally, we further investigated the chosen candidates, prioritizing the peptides with source proteins that have transcript level high fold change for both DESeq analyses and simultaneously a strong binding affinity for both H2K^d^ and H2D^d^ allotypes (cut off values −log_10_ 0.5 H_Average ranks and third quartile of average fold change Fig.4C), generating the final list of candidates (Table 1).

**Figure 4.**
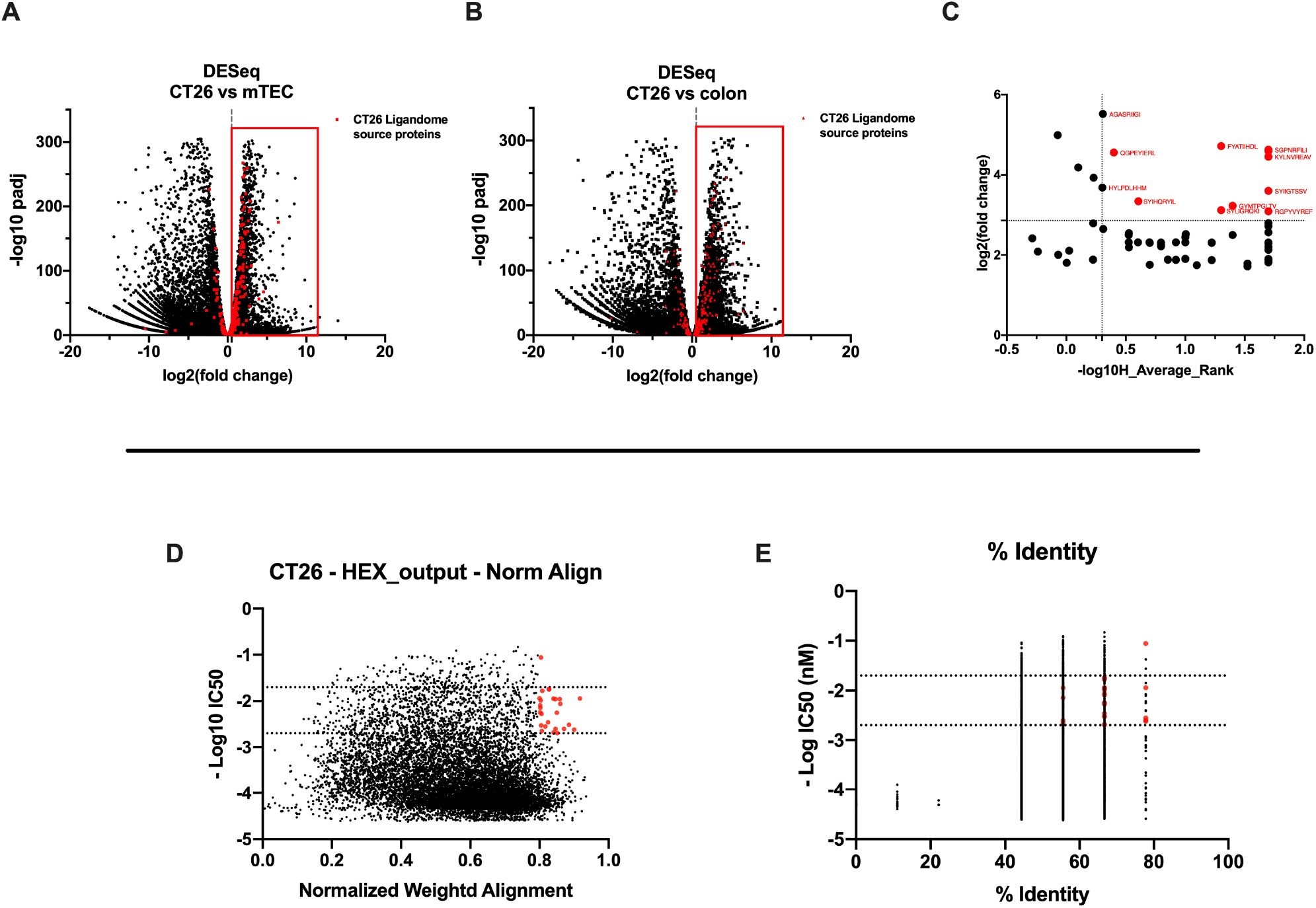
Differential expression and HEX analysis for the MHC-I ligand candidates. **A-B)** Differential gene expression profile (DESeq) in CT26 versus mTEC **(A)** and CT26 versus healthy Balb/c colon **(B)** is depicted as volcano plot of −log_1**0**_ of p-adjusted values versus log_2_ ratio (fold change). The source proteins of MHC-I ligands from our data set are marked in red and the difference expression is considered significant for a fold-change of 1.5 and a padj-value of 0.05 (red square). **C)** Scatter plot comparing the fold change of the source proteins found statistically overexpressed in both DESeq analysises and the average binding affinity score for both H2K^d^ and H2D^d^ allotypes. The values were considered significant for > −log_10_ 0.5 H_average ranks and for the third quartile of average fold change (red marked). **D-E)** The peptides were stratified based on their binding affinity expressed as −log_10_ and on the weighted score to prioritize similarity between more central amino acids in the peptide **(D)** or on the percentage of similarity to viral peptides **(E)**. Binding affinity < 50nM and weighted score and similarity >0.8 were considered as the threshold to select tumor peptides similar to viral epitopes.

**TABLE 1.**
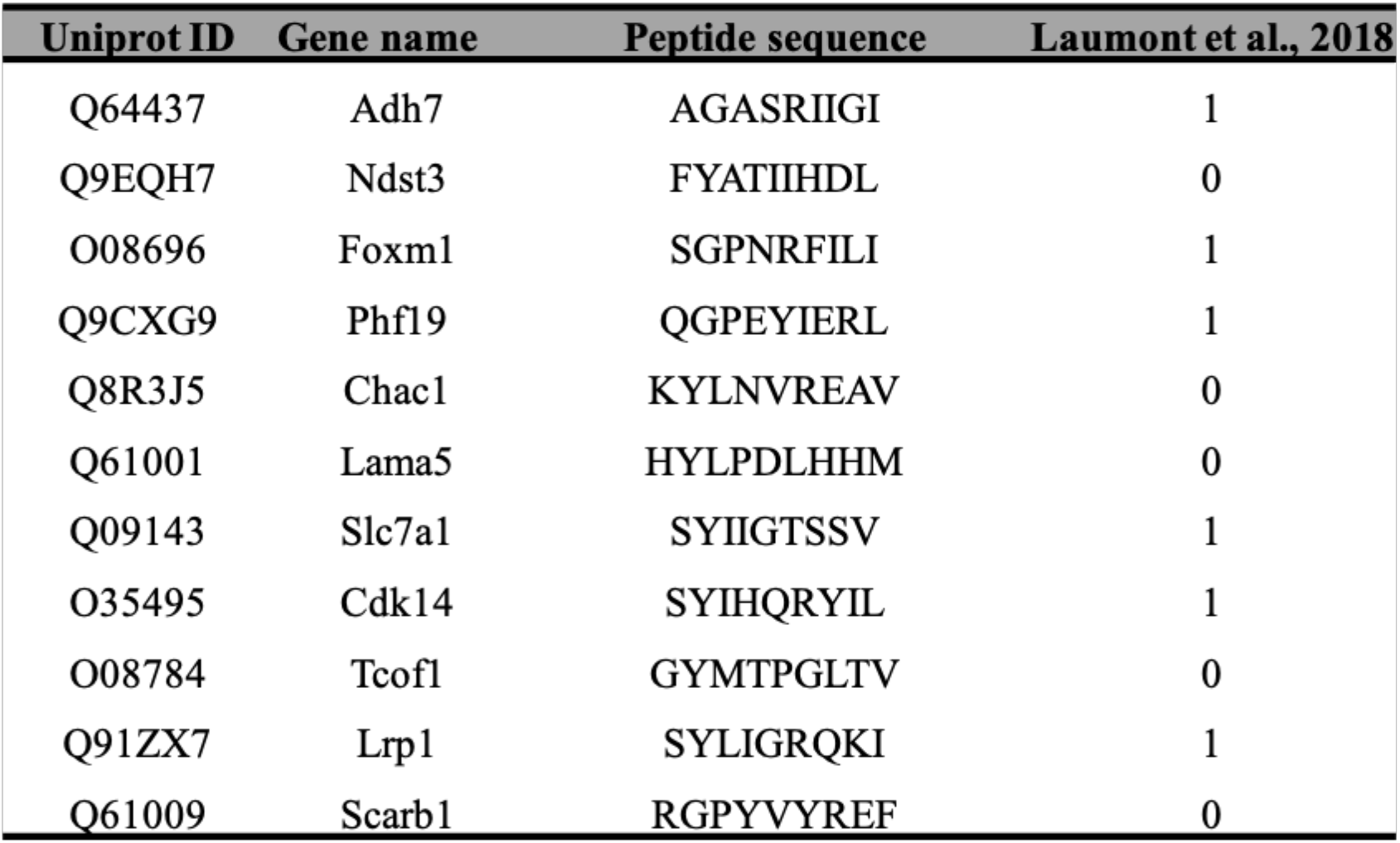
List of candidate peptides derived by differential gene expression profile (DESeq) analysis in CT26 versus mTEC and CT26 versus healthy Balb/c colon. For each peptide, the Uniprot ID, Gene name and the sequence are reported. Additionally, the last column indicates whether (1) or not (0) the peptide has been already described in a published ligandome data set.

The second approach consisted of using the HEX software to inspect the sequences of MHC-I ligands for similarity to antigens from pathogen. First, the software prioritized the peptides that were concurrent strong binders (cut off IC_50_ range 50nM-500nM according to NetMHC4.0) and that showed higher weighted alignment score (cut off 0.8-1 normalized weighted alignment score). The latter focuses on the peptidés similarity in the area of interaction that most likely will engage the TCR of CD8+ T cells, in order to mediate immune response (Fig.4D); the resultant peptides are then further categorized based on their overall percentage of identity to various pathogen antigens and IC_50_ binding affinity score (Fig.4D). The ultimate output consisted in thirteen peptides with their counterpart pathogen peptides (Table 2). Thus, the list of candidates derived from RNAseq analysis and HEX software accounted for 26 peptides. The peptides where then functionally characterized in *in vivo* setting. To determine the peptide immunogenicity, mice were pre-immunized with subcutaneous injection of each peptide in presence of the adjuvant Poly(I:C) and a group of mice was injected either with Poly(I:C) alone or saline as control as well.

**TABLE 2.**
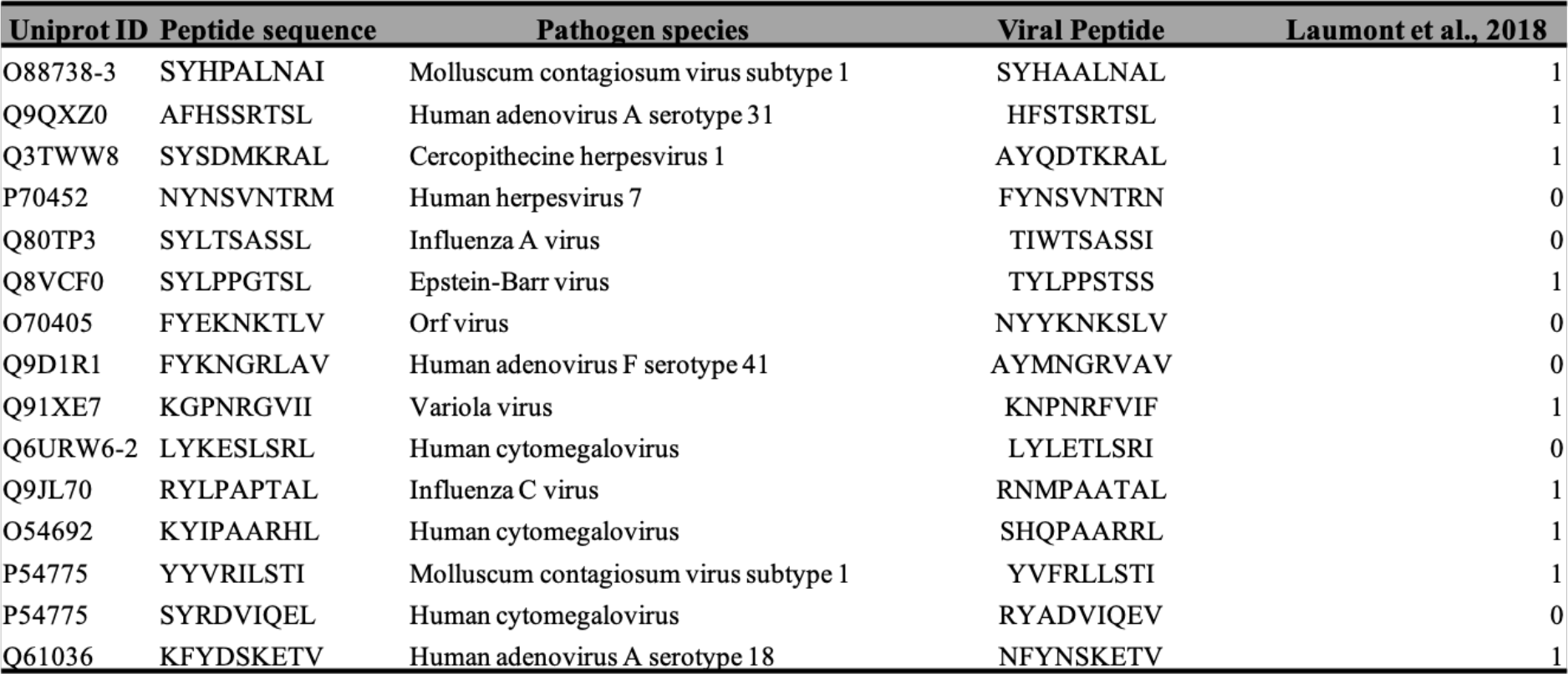
HEX software results. For each peptide, the Uniprot ID, the aminoacid sequence, the similar pathogen species with the respective viral peptides with sequence similarity are shown. The last column indicates whether (1) or not (0) the peptide has been already described in a published ligandome data set.

The splenocytes from those mice were harvested and tested for IFNγ production upon specific stimuli in an ELISpot assay, according to the peptide identification number presented in Table 3. Our data showed that six peptides induced higher frequencies of T cell specific response (Fig.5**, red squares**) defined as the average of the number of spots above the threshold of at least one hundred (peptide 4) that is at 10fold change compared to the control groups. Next, the six peptides selected in the ELISpot assay were modified to contain poly-lysine attachment moiety (polyK-peptides) at the N-terminus to increase the net charge at pH7 (Table 4) and tested for their electrostatic interaction with the OAd; to this end, APTES silica SiO_2_ sensors were first coated with the VALO-mD901 and then 100uM of polyK-peptides were injected into the surface plasmon resonance (SPR) system. Peptide 7 is gp70_423–431_ (AH1-5), a known immunodominant antigen of CT26 derived from a viral envelope glycoprotein encoded in the genome and it was analyzed as well to exploit it as control in downstream animal experiment. The interactions of OAd with the peptides were measured at equilibrium (MAX) and at dissociation (MIN) points (Fig.6A). At equilibrium point, all peptides showed interactions with OAd (Fig.6B**-C)**. However, at dissociation stage, peptide 1, peptide 2, peptide 6 and peptide 7 reached the highest number of peptides retained for viral particle.

**Figure 5.**
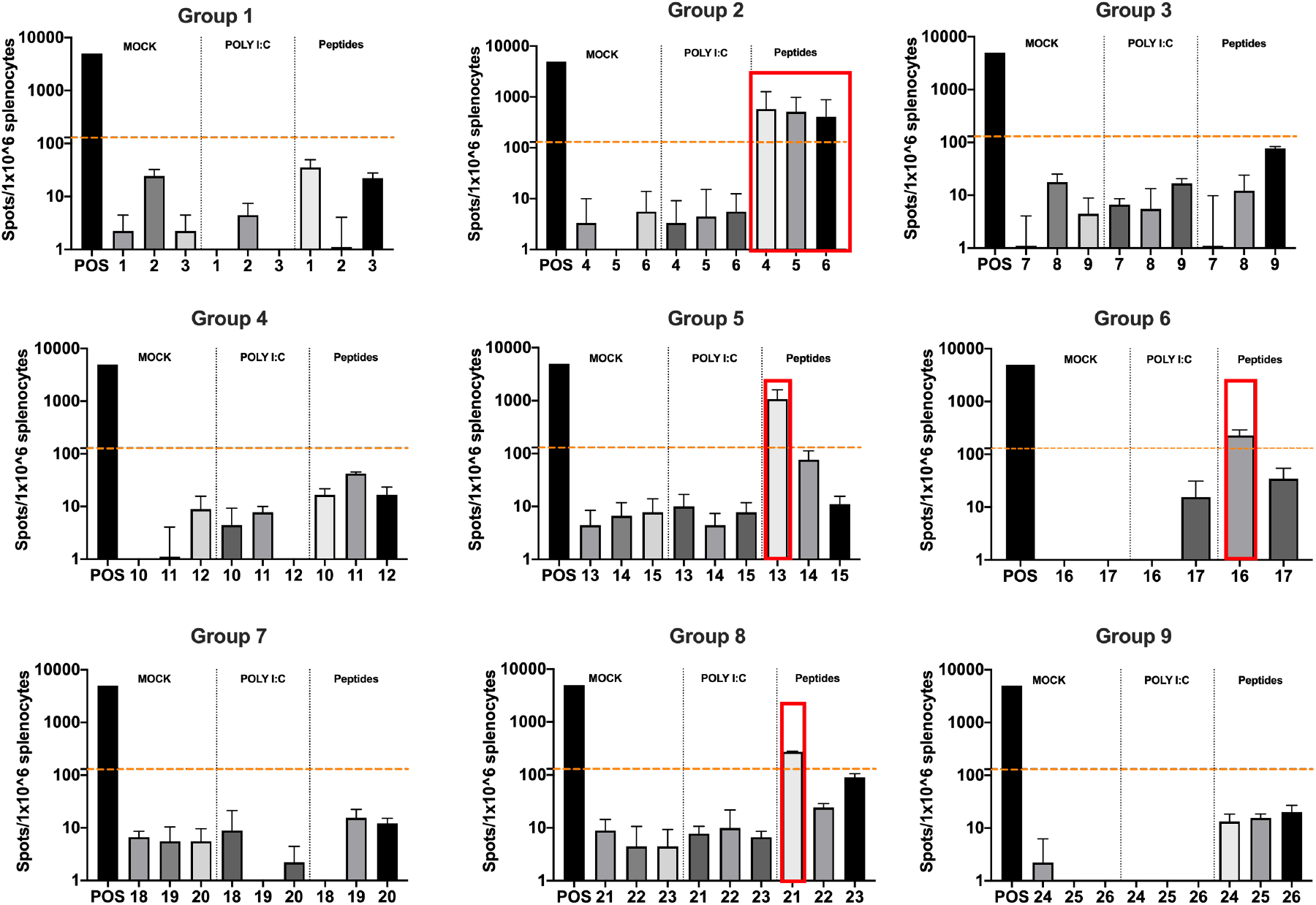
Functional characterization of the peptide candidates. ELISpot IFN-γ analysis was performed on splenocytes harvested from mice pre-immunized with Poly(I:C) and the peptide candidates. The figure shows the stimuli conditions and the treatment groups. The frequencies of anti-tumor T cells responses are depicted as peptides specific reaction per 1×10^6^ splenocytes. The average of the number of spots above one hundred (that is 10fold change compared to the control groupś signal, orange dashed line) was defined as the inclusion criteria to select the peptides (red square).

**Figure 6.**
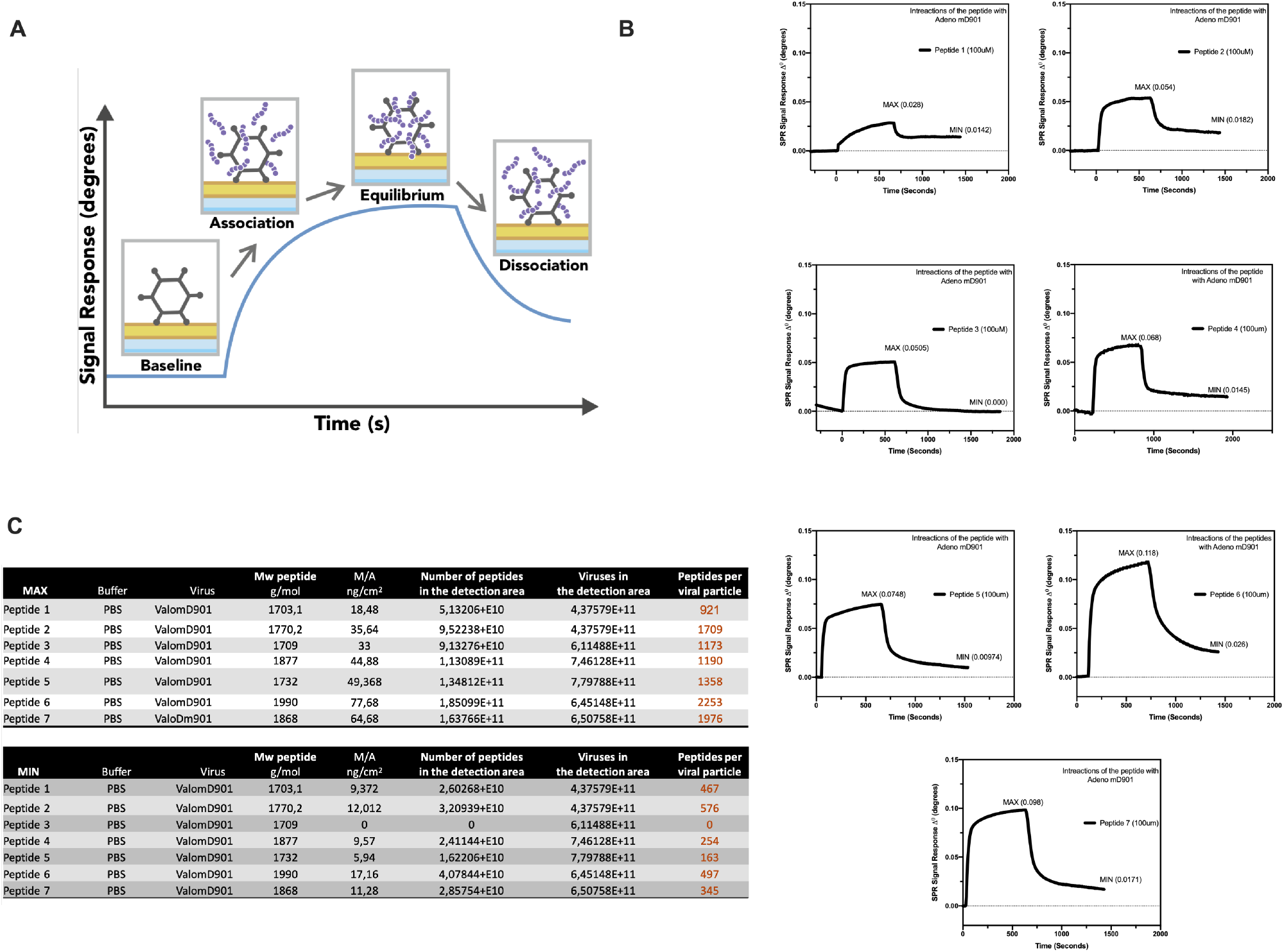
Surface plasmon resonance (SPR) analysis of the peptide/OAd interaction. **A)** An overview of the SPR analysis principle is depicted. **B)** Surface plasmon resonance analysis of the interaction between the poly-lysine modified peptides and OAd is shown as Signal Response degree and time (seconds). For each peptide, the maximum interaction (MAX, equilibrium) and the minimum (MIN, dissociation) peak is reported. **C)** For each peptide and for both equilibrium and dissociation stage, the number of peptides per viral particle has been determined.

**TABLE 3.**
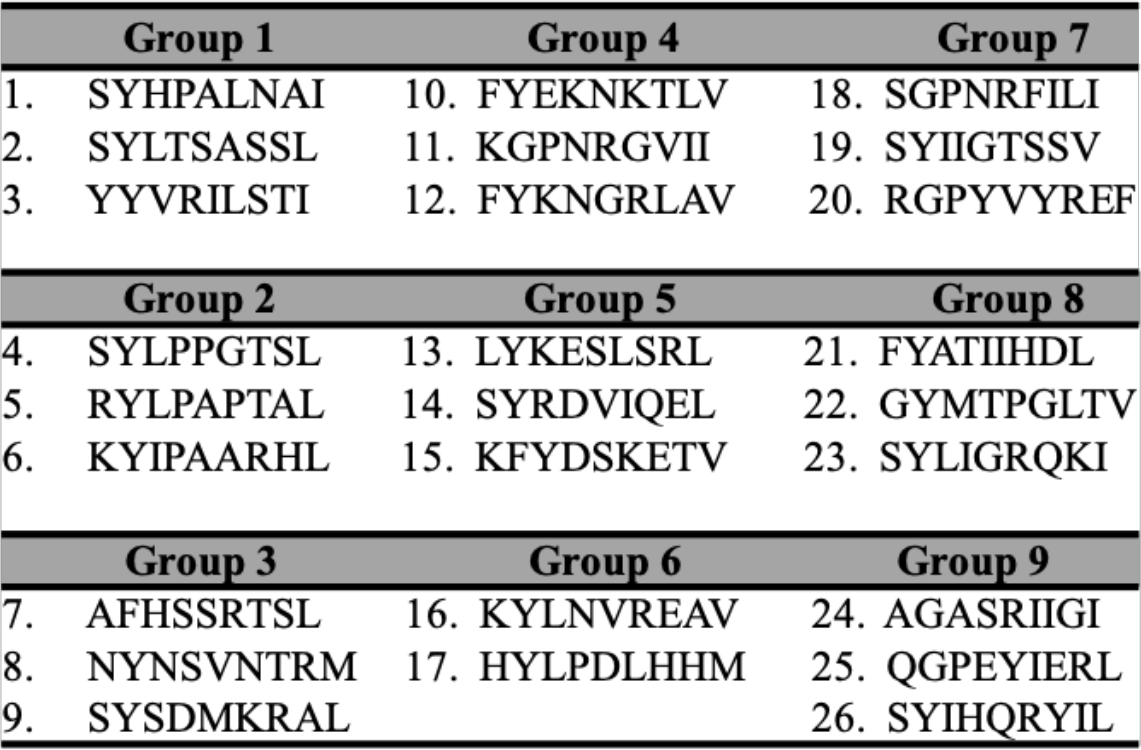
For each group of mice, the peptides with the respective identification number as indicated in the ELISPOT assay is reported.

**TABLE 4.**
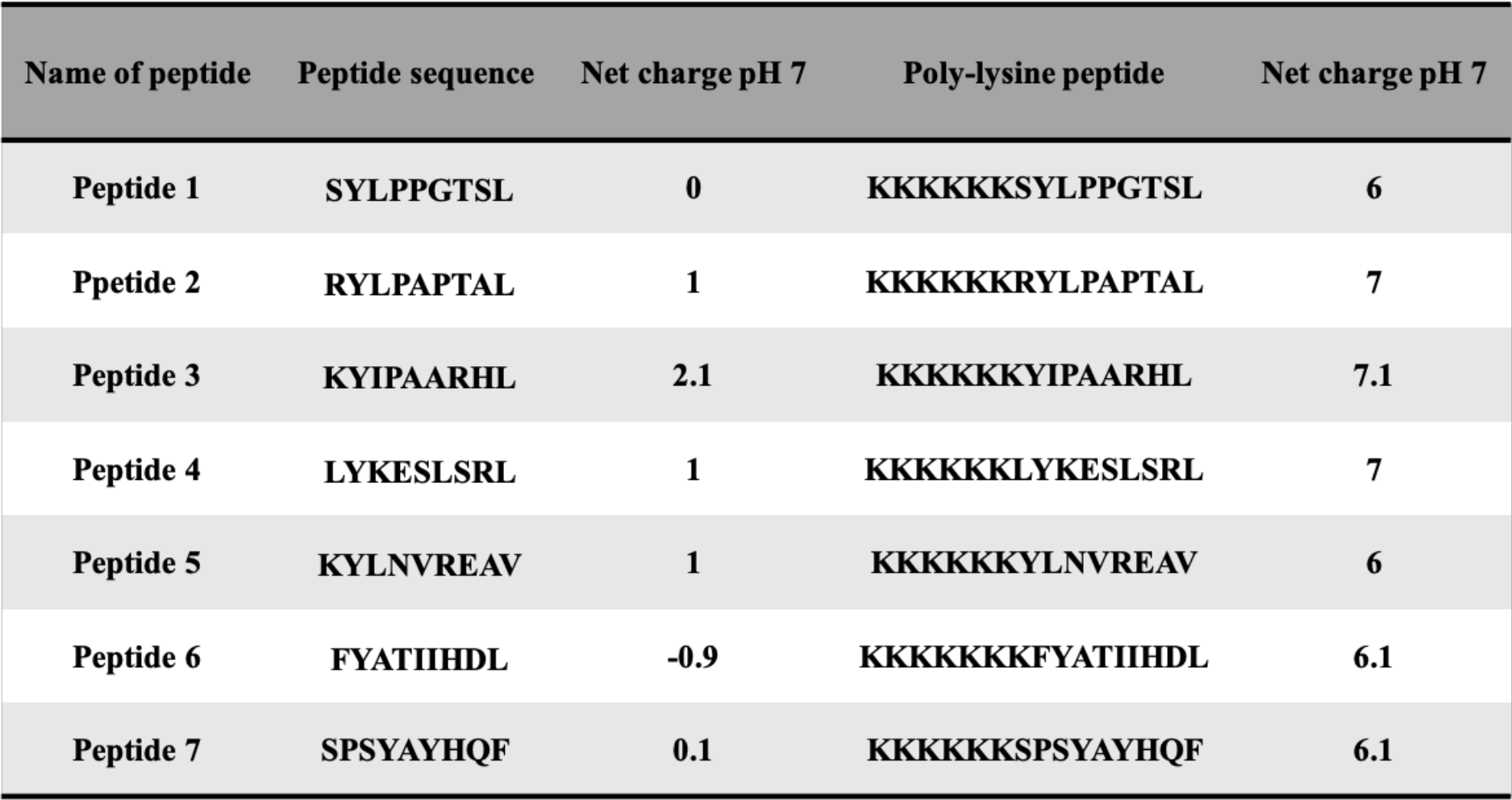
The candidates peptides used in PeptiCRAd technology with the respective net charge without and with the poly-lysine modification is shown.

**TABLE 5.**
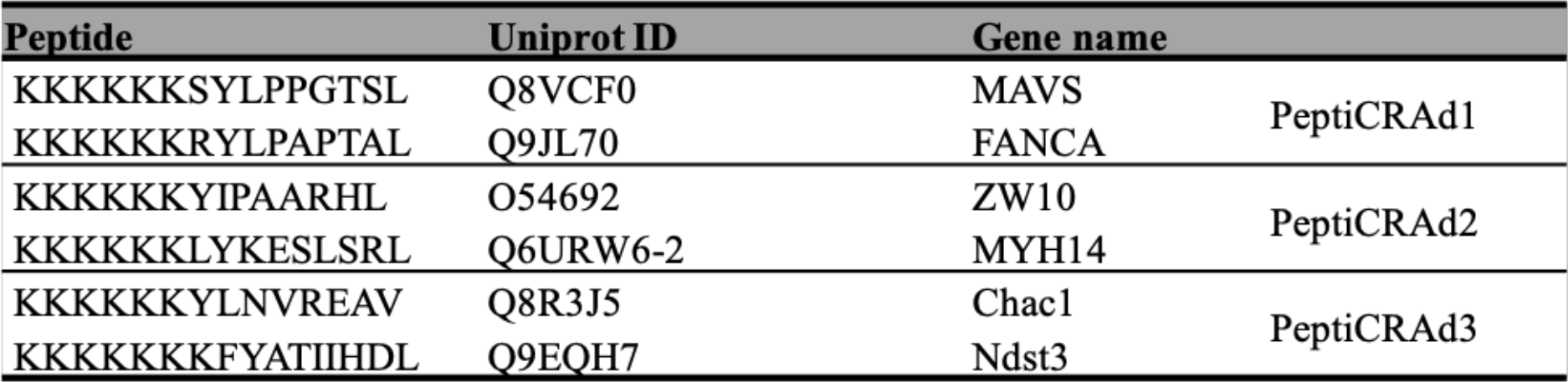
The poly-lysine modified peptides, the Uniprot ID and the respective gene name for each PeptiCRAd treatment group is summarized.

In summary, the *in vitro* and *in vivo* validation and characterization guided the selection of candidate peptides to be used with our PeptiCRAd technology to elicit anti-tumor T cell response.

### PeptiCRAd platform induces systemic anti-tumor immune response controlling the tumor growth of distant untreated cancer lesion in murine model of colon carcinoma

By applying RNA seq and HEX software followed by an *in vivo* functional characterization, we identified six peptides to be tested (Table 4, Peptide 1-Peptide 6) in the PeptiCRAd cancer vaccine platform. The adenovirus used in the PeptiCRAd platform was VALO-mD901, genetically modified to express murine OX40L and CD40L and previously shown to elicit tumor growth control and systemic antitumor response in murine model of melanoma (9). Therefore, immunocompetent Balb/c mice were subcutaneously injected with the syngeneic CT26 tumor cells in the left and right flank. (**Day 0,** Fig.7A). When the tumors were established (**Day 7,** Fig.7A), VALO-mD901 was coated with a pair of each polyK-peptide in our list (PeptiCRAd1, PeptiCRAd2, PeptiCRAd3, Table 4) and injected intratumorally only in the right tumor. PeptiCRAd4 consisted of VALO-mD901 coated with gp70_423–431_ (AH1-5); Mock and VALO-mD901 groups were used as controls. PeptiCRAd1 and PeptiCRAd2 improved tumor growth control as well as VALO-mD901 in the injected lesions (Fig.7B, right panel) as depicted also in the single tumor growth curves per each mouse per each treatment group (Supp.Fig.4). In addition, PeptiCRAd1 (PC1) showed a clear trend towards an improved anti-tumor growth control in the untreated tumor in contrast to all other groups (Fig.7B, left panel). As we sought to investigate the immunological modulation due to the treatments, tumors were harvested for downstream flow cytometric analysis. Interestingly, PeptiCRAd1 showed higher CD8+/CD4+ T cell ratio (Fig. 8A) within the TME of the treated tumor (right side) well in line with an increased CD8+T cell infiltration (Fig. 8B) in both treated (right side) and untreated (left side) cancer lesions. Moreover, the improved tumor growth control achieved in PeptiCRAd1 group correlated with the upregulation of the migratory marker CXCR4 in the CD8+T cell population in both treated and untreated tumors (Fig.8C) and upregulation of effector marker CXCR3 in the CD8+T cell population in the treated lesions (Fig.8D). Exhaustion markers PD1 and TIM3 were also analyzed. The expression of PD1 in CD8+ T cells population showed a tendency to be upregulated in both treated and untreated cancer lesions (Suppl. Fig.5A), suggesting the presence of antigen experienced T cells response. On the other hand, exhausted CD8+T cells phenotypically defined as PD1+ and TIM3+ were downregulated in the untreated lesions; the same tendency was also seen in the treated tumors (Suppl. Fig 5B). We further investigated the CD4+T cell compartment. Our oncolytic cancer vaccine treatment induced a modest downregulation of the CD4+ T cells in both treated and untreated tumors (Suppl. Fig.5C) in line with the increase of CD8+ T cells as aforementioned. The CD4+ population showed upregulation of CXCR4 in the treated tumors in PeptiCRAd1, PeptiCRAd2, PeptiCRAd3 compared to the VALO-mD901-treated tumors; however, no differences were observed when compared to the mock group. Even though the effector marker CXCR3 was downregulated in the untreated and treated tumors, PeptiCRAd1 showed the tendency in upregulating CXCR3 in the untreated lesion (Suppl. Fig.5C). No statistical differences were observed as regard to the antigen experienced or exhausted phenotypes compared to the control groups (Suppl. Fig.5C).

**Figure 7.**
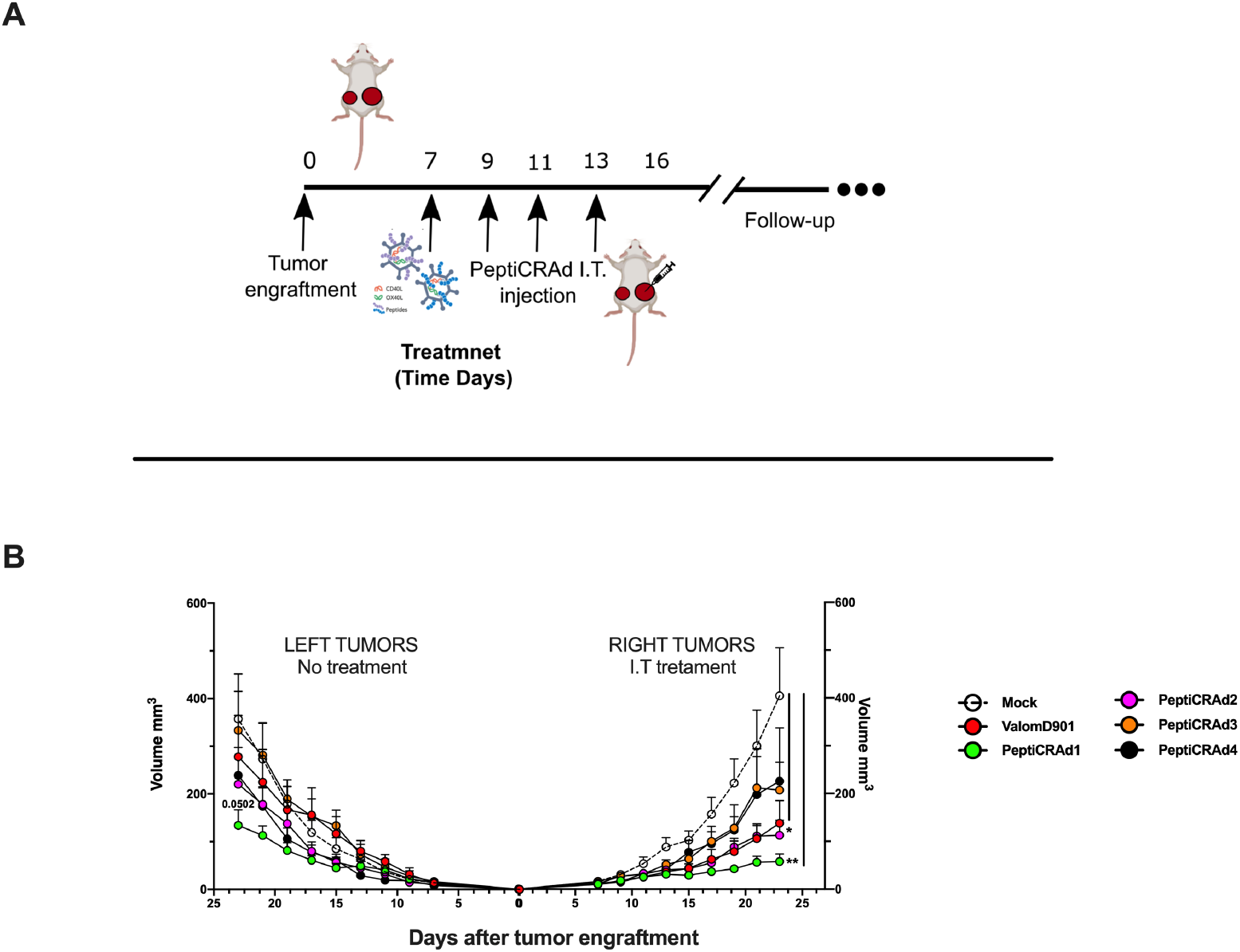
PeptiCRAd improved the tumor growth control in both injected and not injected lesions. **A)** A schematic representation of the animal experiment setting is depicted. Immunocompetent Balb/c mice were subcutaneously injected with the syngeneic tumor model CT26 in the left (0.6×10^6^ cells) and right flank (1×10^6^). PeptiCRAd was intratumorally administrated four times, two days apart. **B)** The CT26 tumor growth was followed until the end of the experiment and the tumor size is presented as the mean ± SEM and statistically difference was assessed with two-way ANOVA; (*, P < 0.05; ***, P < 0.001; ****, P < 0.0001; ns, nonsignificant).

**Figure 8.**
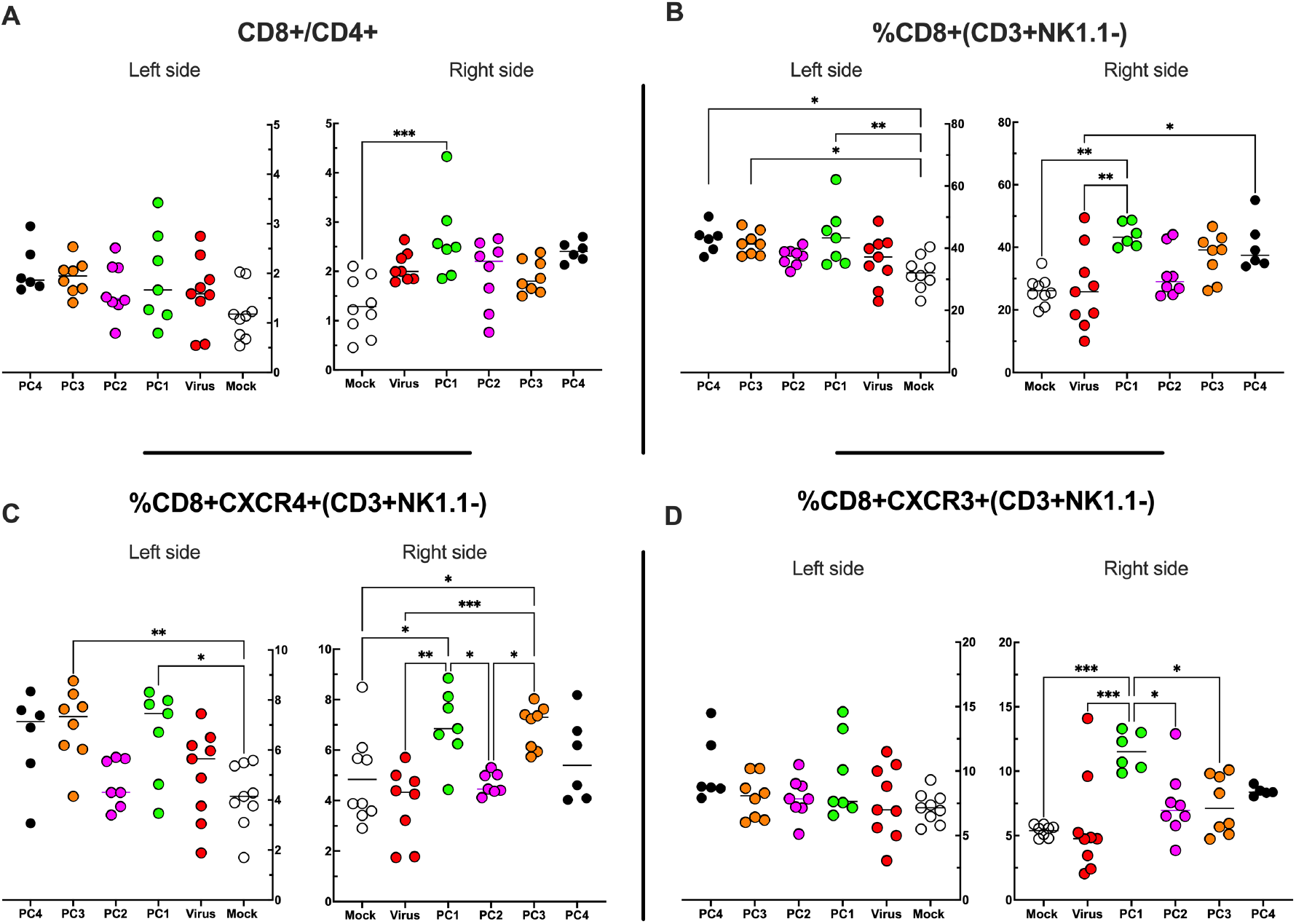
Flow cytometry analysis of Tumor Infiltrating Lymphocytes (TILs). **A-D)** The treated (right side) and the untreated tumors (left side) were harvested at the end of the experiment and analyzed for the CD8+/CD4+ ratio **(A)** and for the frequency of CD8+ **(B)**, CD8+CXCR4+ **(C)**, CD8+CXCR3+**(D)** in the TME. All the data are plotted as dot plot for each mouse and for each treatment group. The significance was assessed by One way ANOVA and Tukeýs correction (*, P < 0.05; ***, P < 0.001; ****, P < 0.0001; ns, nonsignificant).

Altogether, the data showed that PeptiCRAd 1 induced remodulation of the immune cell infiltration within the TME, in particular influencing the CD8+ T cell population.

In conclusion, the pipeline reported herein could considerably facilitate the identification, the prioritization, and the selection of suitable peptide candidates for cancer vaccine. Moreover, we also proposed an easy and fast adenovirus-based platform for the generation of personalized oncolytic vaccines to be combined with the selected peptides for cancer immunotherapeutic treatments. We envision that our pipeline could be applied to human clinical approaches, drastically reducing the time related to both tumor peptide selection and oncolytic vaccine generation, paving the way to precision cancer immunotherapy treatments.

## DISCUSSION

Cytotoxic anti-tumor CD8+ T cells (CTLs) recognize peptides typically of 8-10 amino acids within the MHC-I complex expressed on the cellular surface and therefore the knowledge of these peptides is the key to design T cell based therapeutic cancer vaccines; indeed, their efficacy relies mostly on the choice of the antigenic peptides (21). These peptides should be highly immunogenic, expressed exclusively on the cancer cells to avoid on-target off-tumor toxicity and tailored on the patient’s specific tumor ligandome landscape. However, only a fewer if any of the tumor antigens meet those characteristics, making it very difficult to generate peptide-based vaccination technologies. Thus, the isolation and identification of MHC-I peptides and the subsequent selection criteria are of utmost importance in creating those vaccines. To fulfill these needs, we conceived a pipeline that comprises all the steps considered essential for an optimal development of a therapeutic cancer vaccine.

We decided to identify and isolate peptides directly from the MHC-I complexes, exploiting state-of-the-art immunoprecipitation and mass spectrometric methodologies as the direct elution and analysis of MHC-I restricted peptides is so far the most reliable and used approach in studies of ligandome landscape (2), identifying naturally processed and presented tumor epitopes that could generate clinically relevant anti-tumor responses. Even tough computational algorithms can take into account the entire MHC complex presentation machinery (e.g., proteasomal cleavage, transporter-associated antigen processing (TAP) transport, binding motif) to predict relevant T cell epitopes, the lack of validated and homogenous datasets makes the process difficult and less reproducible (22, 23). These considerations prompted us to adopt direct MHC-I immunoaffinity purification as first step of our pipeline.

Moreover, to develop and further validate our proof-of-concept pipeline, the choice of the tumor model needed to meet specific requirements. First, we wanted a tumor model that expresses sufficient levels of MHC-I complexes, granting a fruitful recovery of peptides from the cellular surface. Indeed, the overall idea was to obtain a conspicuous list of peptides, in order to later on challenge our prioritizing and selection criteria. Secondly, to test the selected candidates for their anti-tumoral efficacy profile, the pre-clinical model should have beneficial immunogenic features, in particular T cell infiltration into the TME, allowing a better study of the immune modulation upon treatment administration. Based on that, the colon tumor model CT26 was selected as it showed high expression level of MHC-I complexes as demonstrated in our flow cytometry analysis and for being a widely used and characterized tumor model for developing and testing immunotherapeutic concepts *in vivo* (24, 25). As expected, the immunoaffinity purification generated a long list of peptides, containing more than 8000. Before moving forward in our pipeline, we carefully analyzed the quality of the produced data set to ensure the solidity of our list and to examinate it for the presence of contaminants. The analysis demonstrated that the eluted peptides resembled a typical ligandome profile and therefore they could be considered as true MHC-I ligands. The strength and the reliability of the ligandome data set is critically important for the following steps as it influences the subsequent results.

Of note, beside the identification of MHC-I peptides, the main issue is dealing with the prioritization of the peptides among thousands of possible candidates. In this context, we followed two parallel directions. First, we adopted a more conservative approach that consisted in analyzing the RNAseq expression level of the respective source proteins. In particular, based on the definition of TAA as an antigen overexpressed in malignant cells compared to healthy tissue, we considered the colon Balb/c as the reference normal tissue since CT26 is an undifferentiated colon carcinoma induced by the carcinogen N-nitroso-N-methylurethane (NMU) (25). In this sense, the selected peptides used as therapeutic cancer vaccine should evoke specific anti-tumor CTLs able to recognize and eradicate tumor cells, avoiding damages to normal colon tissue. Moreover, syngeneic mTEC (medullary thymic epithelial cells) expresses most of the known genes and it is the site of T cell selection to induce central tolerance to MHC peptides coded by their vast transcriptome. We assumed that the breaking of tolerance could most likely happen if the source proteins of the selected peptides from our data set were overrepresented in the CT26 cell line compared to the mTEC. To ensure a more accurate selection of the candidates, we focused the choice on peptides that meet both criteria of I) source protein overexpressed compared to normal colon and mTEC and II) of being true MHC-I strong ligands. The second parallel approach represents the main novelty introduced in our pipeline and it consisted in selecting peptides based on their similarity to antigen pathogen by exploiting HEX, a tool previously developed in our laboratory and successfully validated both in pre-clinical and clinical settings (1).The main idea relies on the intrinsic degeneracy of the T cell receptor (TCR), defined as the ability of a single TCR to recognize more than one antigen, generating a phenomenon known as cross-reactivity. This property is an essential feature to broaden the breadth of the T cell repertoire and for instance it allows anti-viral memory CD8+T cells generated by prior infections to recognize unrelated viruses, as demonstrated in several studies in human and murine models (26, 27). We thought that the same concept could be applied to cancer antigens that have similarities to viral antigens. We are aware that in this work we used mice naïve to viral infections and therefore no memory CD8+ T cell cross-reactivity could be exploited. However, translated in a real clinical setting, our approach will have the added value of exploiting the cross-reactivity of pre-existing viral CD8+T cells to enhance the anti-tumor response. Applying the aforementioned *in silico* analysis, the number of candidates was shortened, making it feasible to further functionally characterized the list of the peptides in an ELISpot assay.

After the selection of candidates to exploit in a vaccine platform were selected, we employed the peptides in our previously developed platform named PeptiCRAd, an oncolytic adenovirus coated with polyK modified peptides (7–9, 28). Indeed, after the FDA approval of T-VEC, a herpes virus encoding GM-CSF (29) for the treatment of melanoma, the use of oncolytic viruses has been extensively explored in cancer immunotherapy (30–33). Oncolytic viruses (OVs) are naturally occurring or genetically modified viruses able to infect and replicate in cancer cells; the OVs induce a systematic immune response, involving both innate and adaptive immune response. Moreover, the antigen spread following the viral burst acts as *in situ* cancer vaccine but is often not enough to generate a specific anti-tumor adaptive immune response, instead generate mainly an anti-viral T cell responses (34). To overcome this limitation, we decided to combine the immunogenicity of the oncolytic viruses with the anti-tumor specificity of the peptides, generating an oncolytic cancer vaccine. Thus, to challenge our pipeline and investigate whether our selection criteria could actually be used to identify relevant candidates for cancer treatment, we decorated the OAd VALO-mD901 with the selected peptides to treat immunocompetent CT26-tumor bearing mice. To understand whether our technology could actually evoke a systemic anti-tumor immune response, we engrafted two tumors for each mouse, both right and left flank and then we treated only the tumor on the right flank. Of note, VALO-mD901 is encoding murine CD40L and OX40L under CMV promoter allowing transgene expression in murine cells. Stimulation of innate (due to CD40L) and adaptive (due to OX40L) immune cells explained the local anti-tumor activity in virus-injected tumors observed in our results and the lack of efficacy in the distant lesions. Contrarily, PeptiCRAd 1 (virus coated with peptide 1) treatment slowed the tumor growth of both the treated and untreated lesions, highlighting the generation of a systemic tumor-specific immune response.

The overall results demonstrated the feasibility of applying the described pipeline for the generation of a tailored therapeutic cancer vaccine. We have addressed all the main issues universally recognized as challenges in the field with main focus on the prioritization and selection criteria among thousands of peptide candidates. Additionally, we adapted quick “plug-and-play” technology based on decorating an OV with the selected peptides. The nature of this technology opens the possibility of a fast generation of tailored therapeutic cancer vaccines in future clinical application where personalized therapies represent one of the main goals for a successful treatment. From a clinical application point of view, the integration of the ligandome and transcriptome analysis could benefit from the fast selection of peptides done with the HEX software. Indeed, recently data suggest that MHC-I restricted peptides homologous to viral peptides are strongly immunogenic and offer a reliable source of candidates for cancer vaccine design. Our approach will capitalize on pre-existing cross-reactive T cells (35, 36), facilitating the peptide selection.

## Acknowledgements

We thank all the participants for their support and advice. Moreover, flow cytometry analysis was performed at the HiLife Flow Cytometry Unit, University of Helsinki. This work has been supported by the European Research Council under the European Union’s Horizon 2020 Framework programme (H2020)/ERC-CoG-2015 Grant Agreement No. 681219, the Helsinki Institute of Life Science (HiLIFE), the Jane and Aatos Erkko Foundation (decision 19072019), the Cancer Society of Finland (Syöpäjärjestöt), ERC (POC), Business Finland.

## Supplementary Figures and Supplementary Figure Legends

**Supplementary Figure 1.**
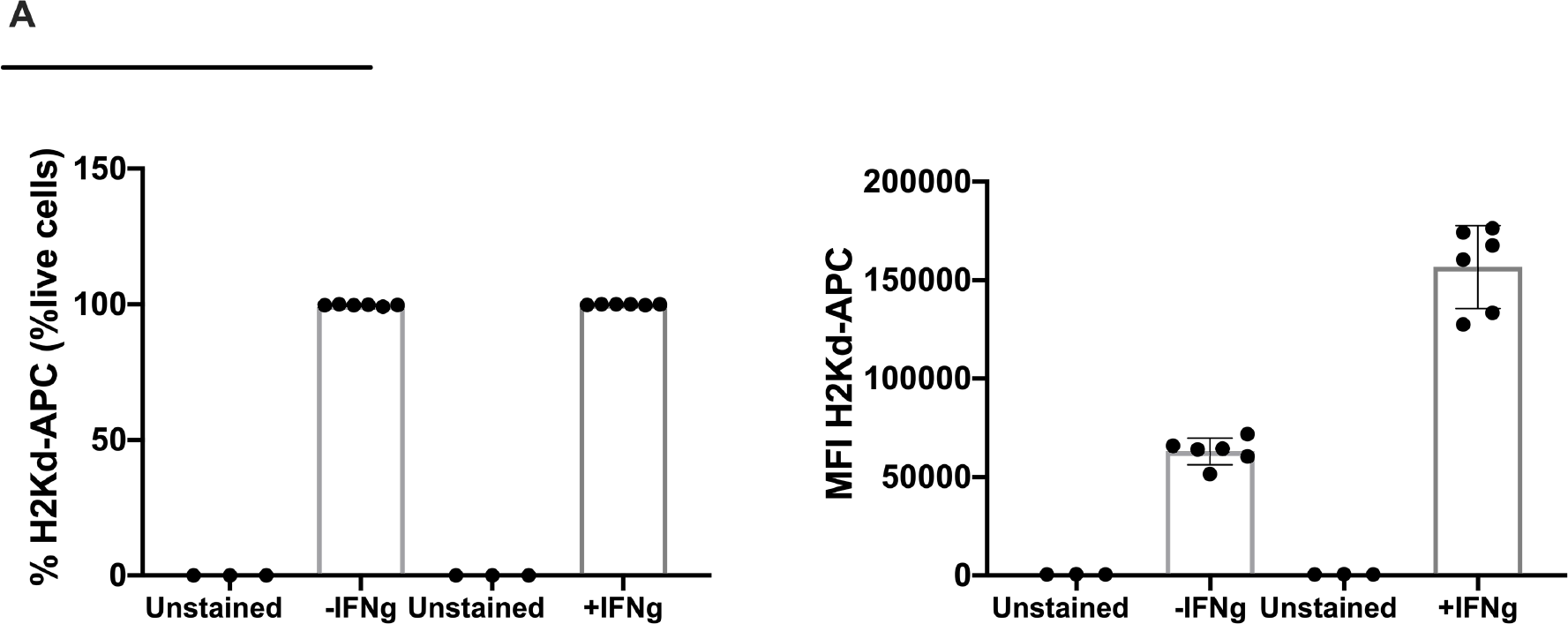
Flow cytometry analysis of H2K^d^ expression level in the colon tumor model CT26. The frequency and the mean fluorescent intensity (MFI) are shown without or upon IFN-γ stimulation.

**Supplementary Figure 2.**
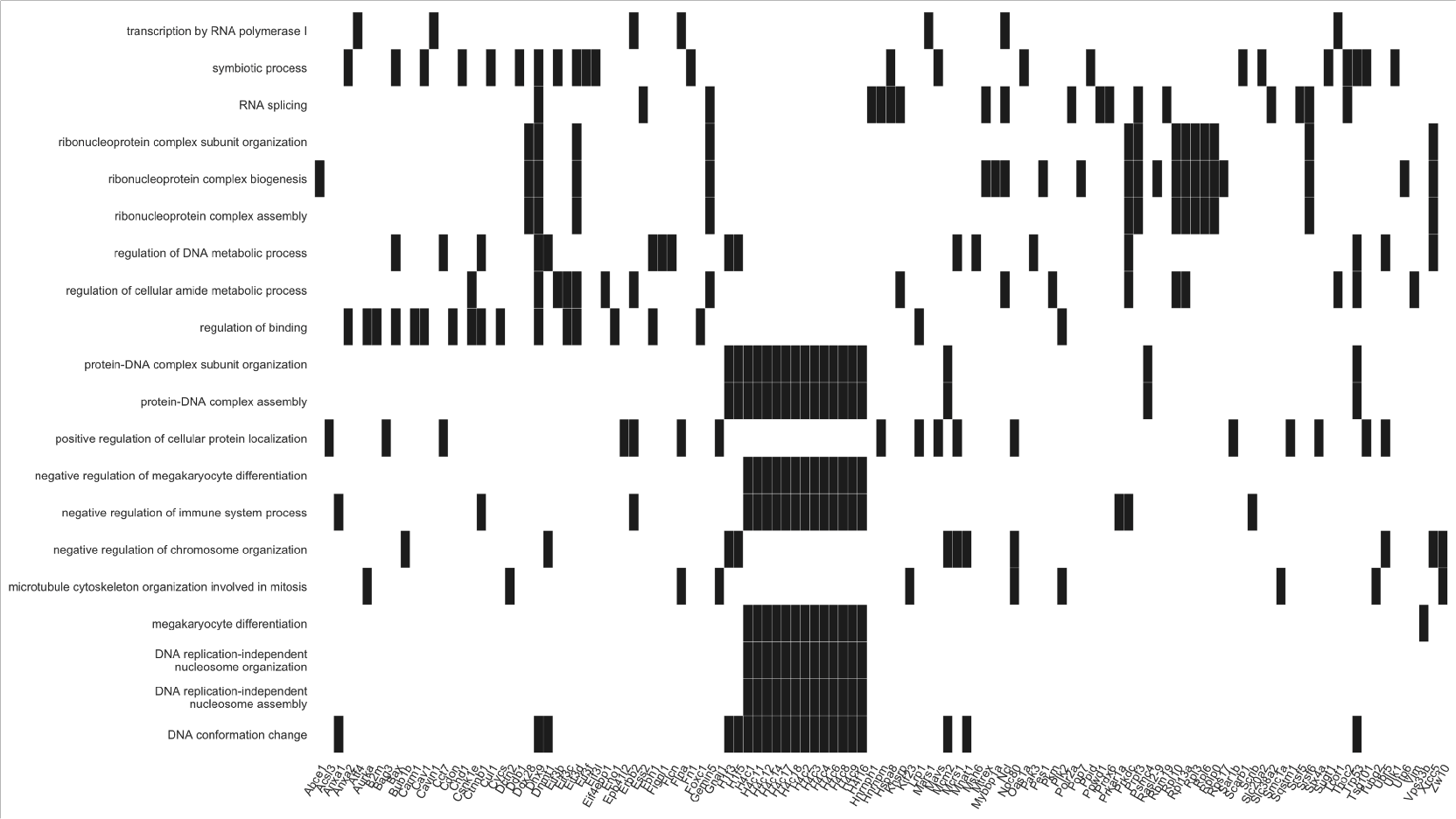
Heatmap of the Gene Ontology (GO) enrichment results are displayed. The biological process analysis of the source proteins was performed and the first twenty biological processes with the respective gene names are shown.

**Supplementary Figure 3.**
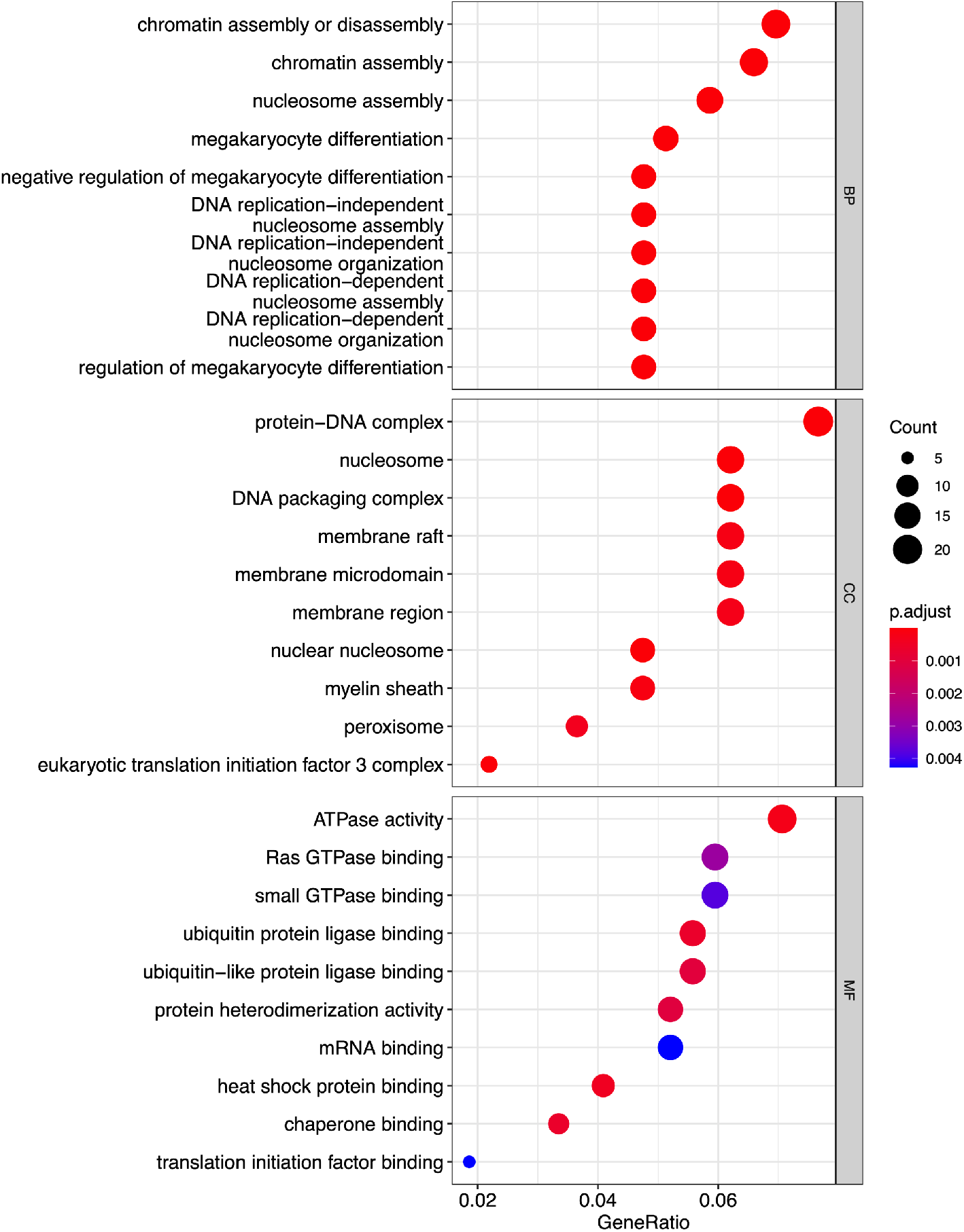
Dot plot showing enrichment of Gene Ontology (GO) biological process (BP), cellular components (CC) and molecular functions (MF); adjusted p-values of the first 10 statically relevant terms are depicted as color gradient.

**Supplementary Figure 4.**
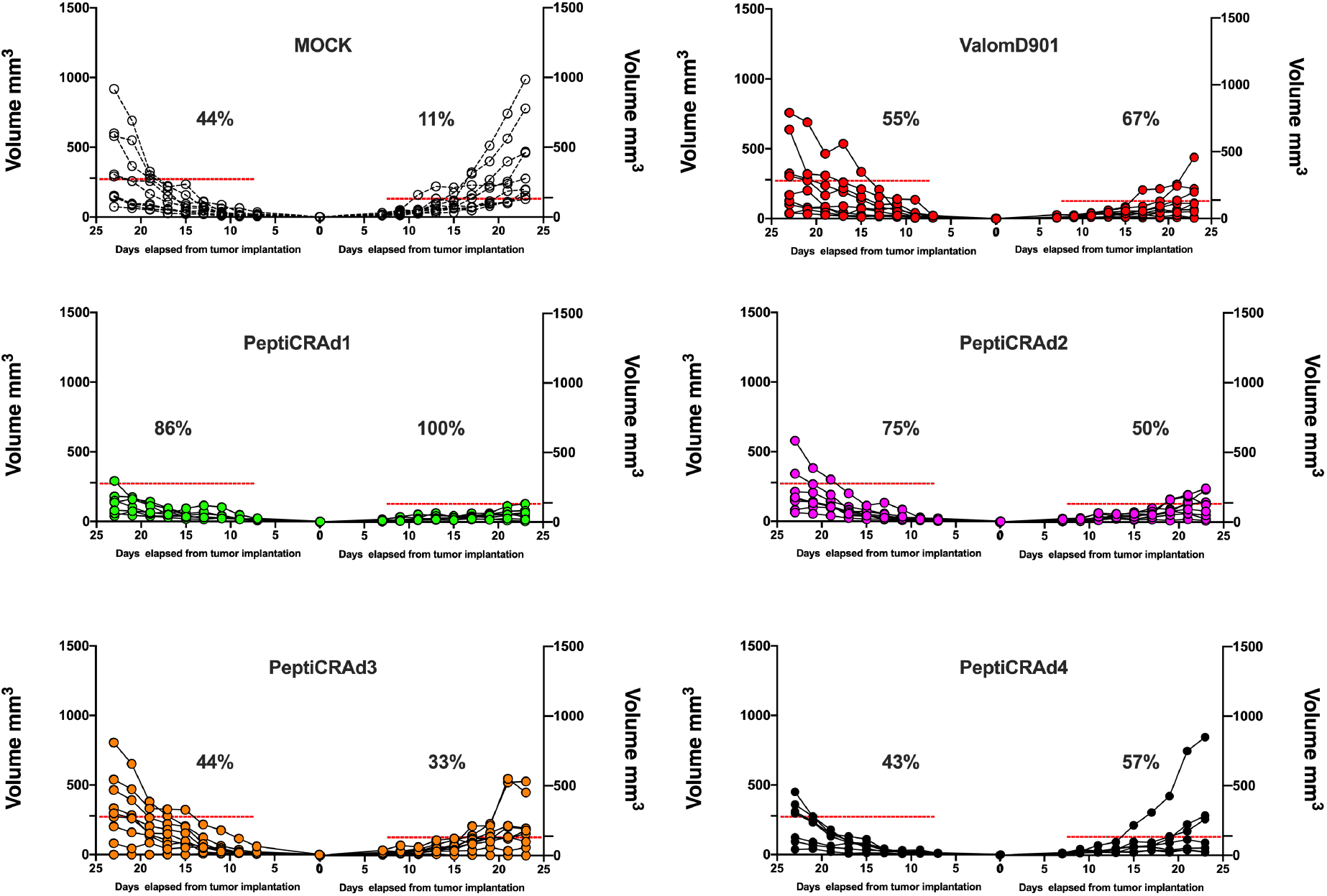
Single tumor growth for single mouse for each treatment group is depicted. A threshold of 138 mm^3^ (right tumor) and 278 mm^3^ (left tumor) was set to define the percentage of mice responding to the different therapies (dotted line). The percentage of responders in each treatment group is shown on the right side of the dotted line. (The threshold was defined as the average of the tumor size at the last day of the experiment in the treatment control group ValomD901and calculated separately for the right and the left tumor).

**Supplementary Figure 5.**
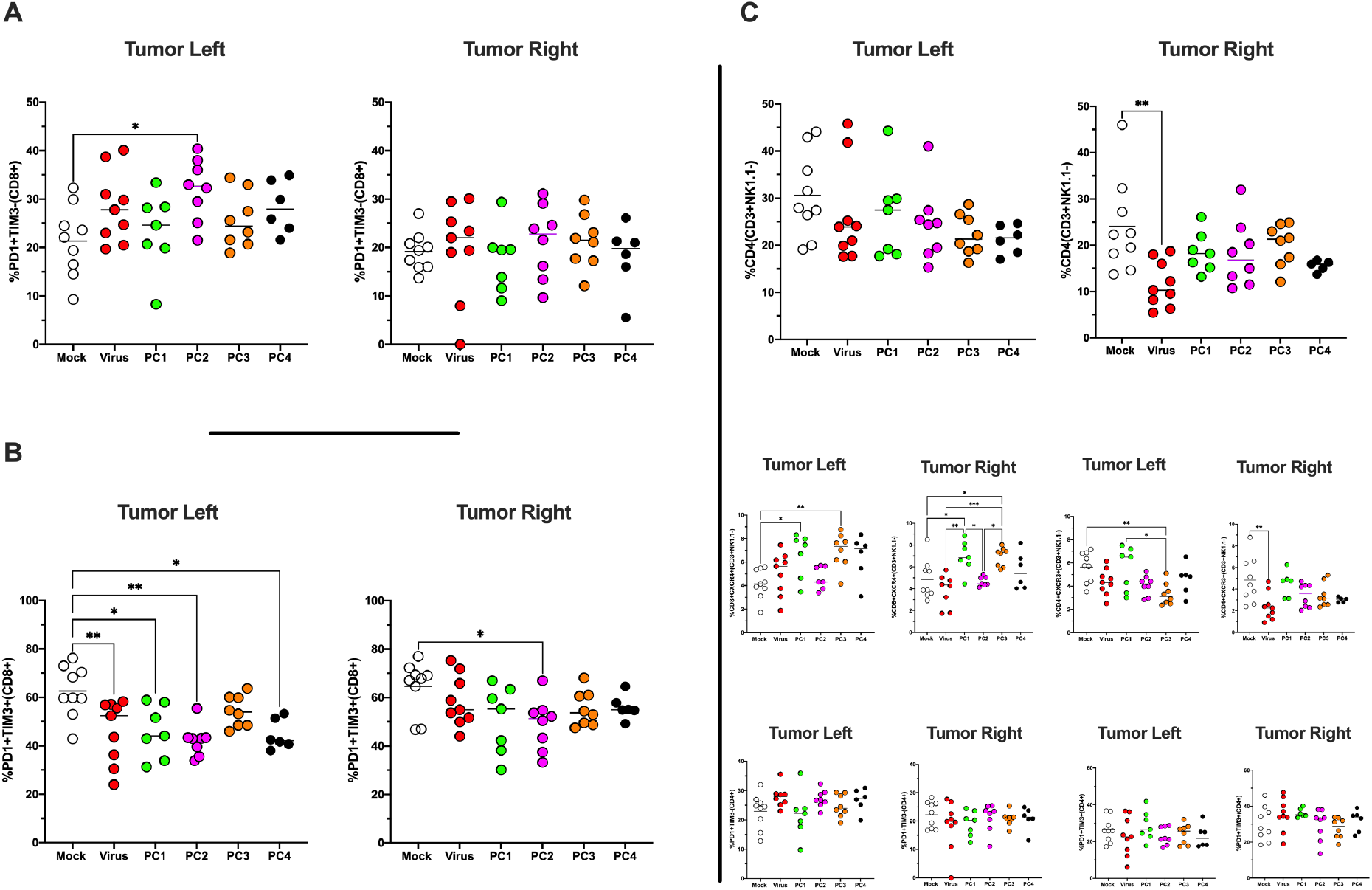
**A-B)** The antigen experience (PD1+TIM3-) **(A)**/exhaustion profile (PD1+TIM3+) **(B)** of CD8+ T cells in the TME was investigated by Flow cytometry analysis **C)** The frequency of CD4+, CD4+CXCR4+, CD4+CXCR3+, CD4+PD1+TIM3-and CD4+PD1+TIM3+ in the TME are shown. All the data are plotted as dot plot for each mouse, for each tumor and for each treatment group. The significance was assessed by One way ANOVA and Tukeýs correction (*, P < 0.05; ***, P < 0.001; ****, P < 0.0001; ns, nonsignificant).

