## Supplementary figures and images for "A novel immunopeptidomic-based pipeline for the generation of personalized oncolytic cancer vaccines"

### Supplementary Figure 1

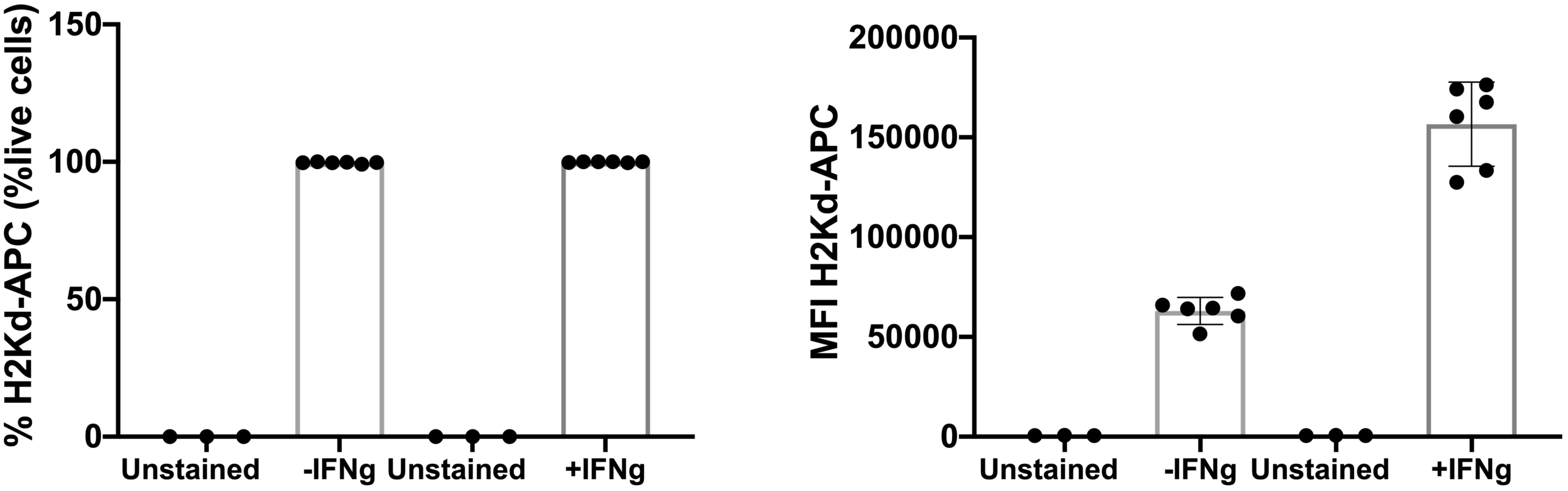

### Supplementary Figure 2

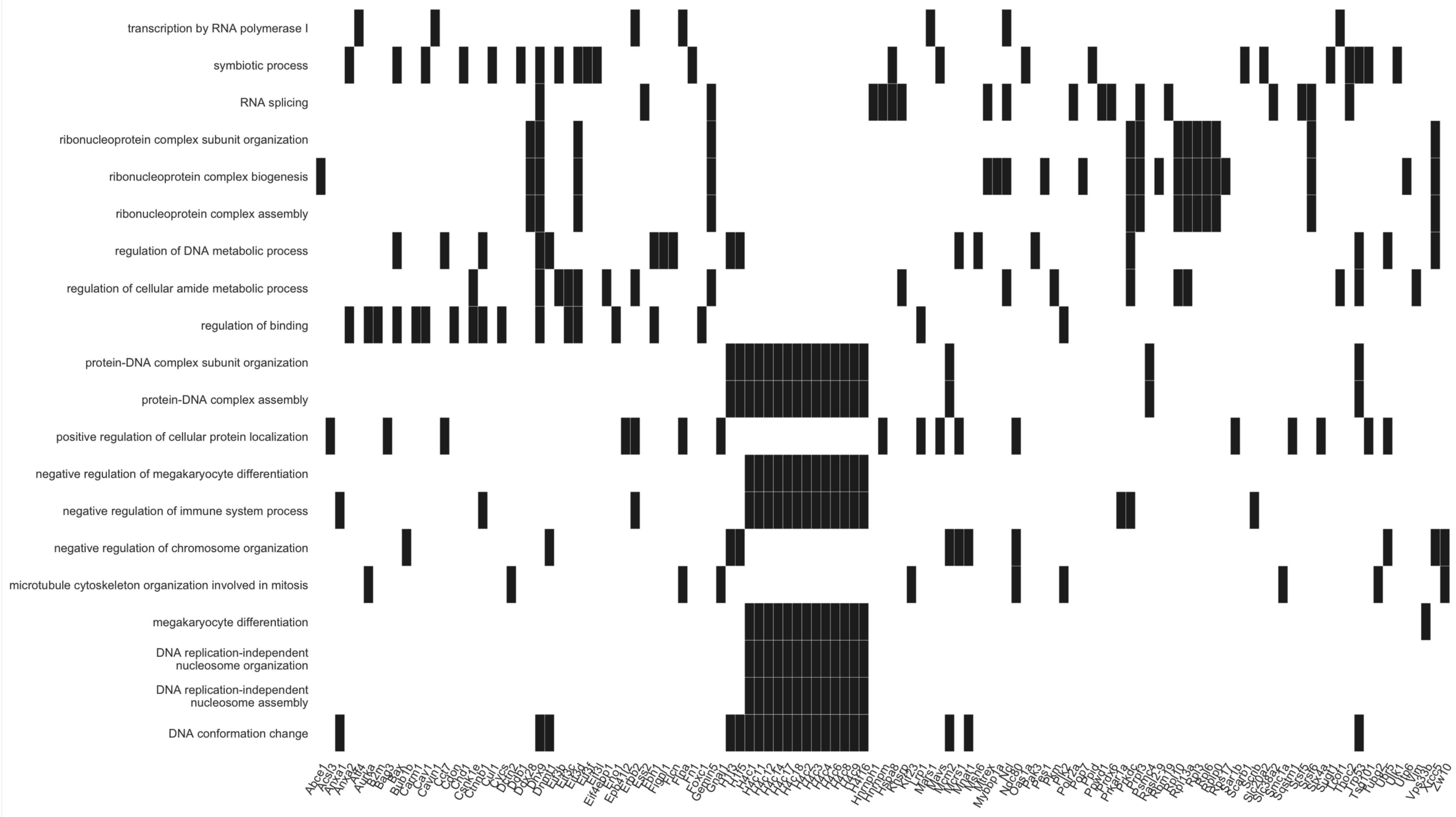

### Supplementary Figure 3

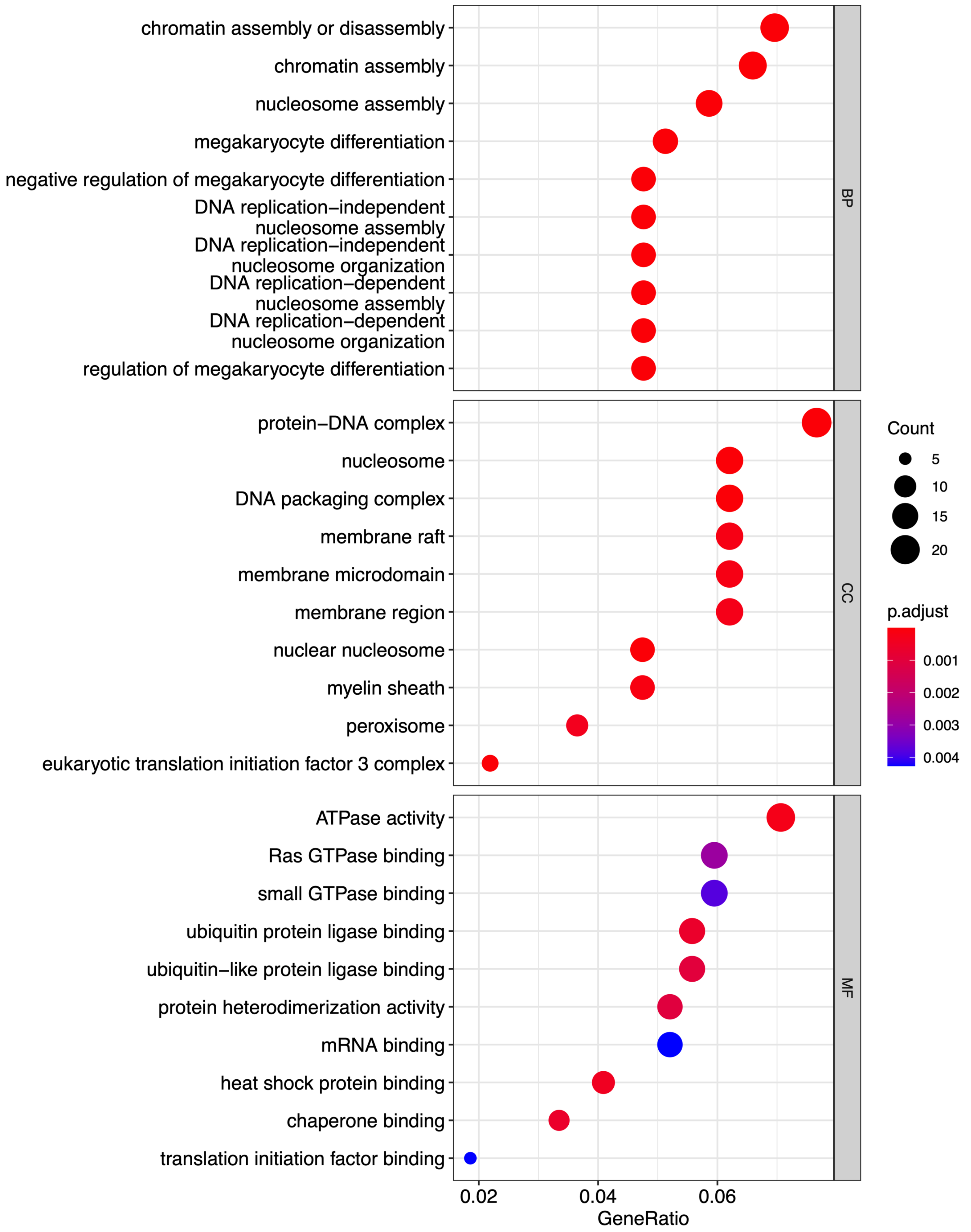

### Supplementary Figure 4

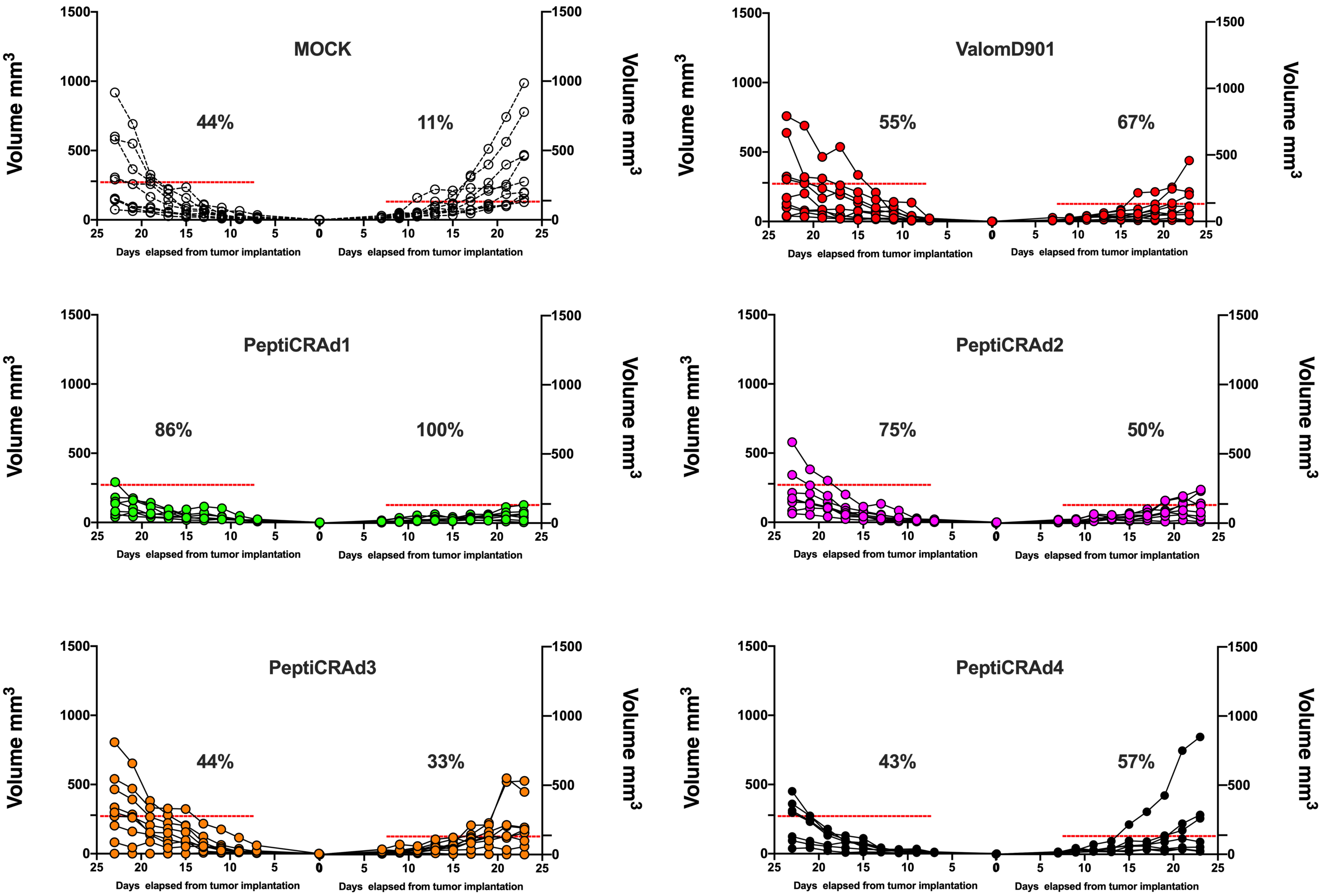

### Supplementary Figure 5

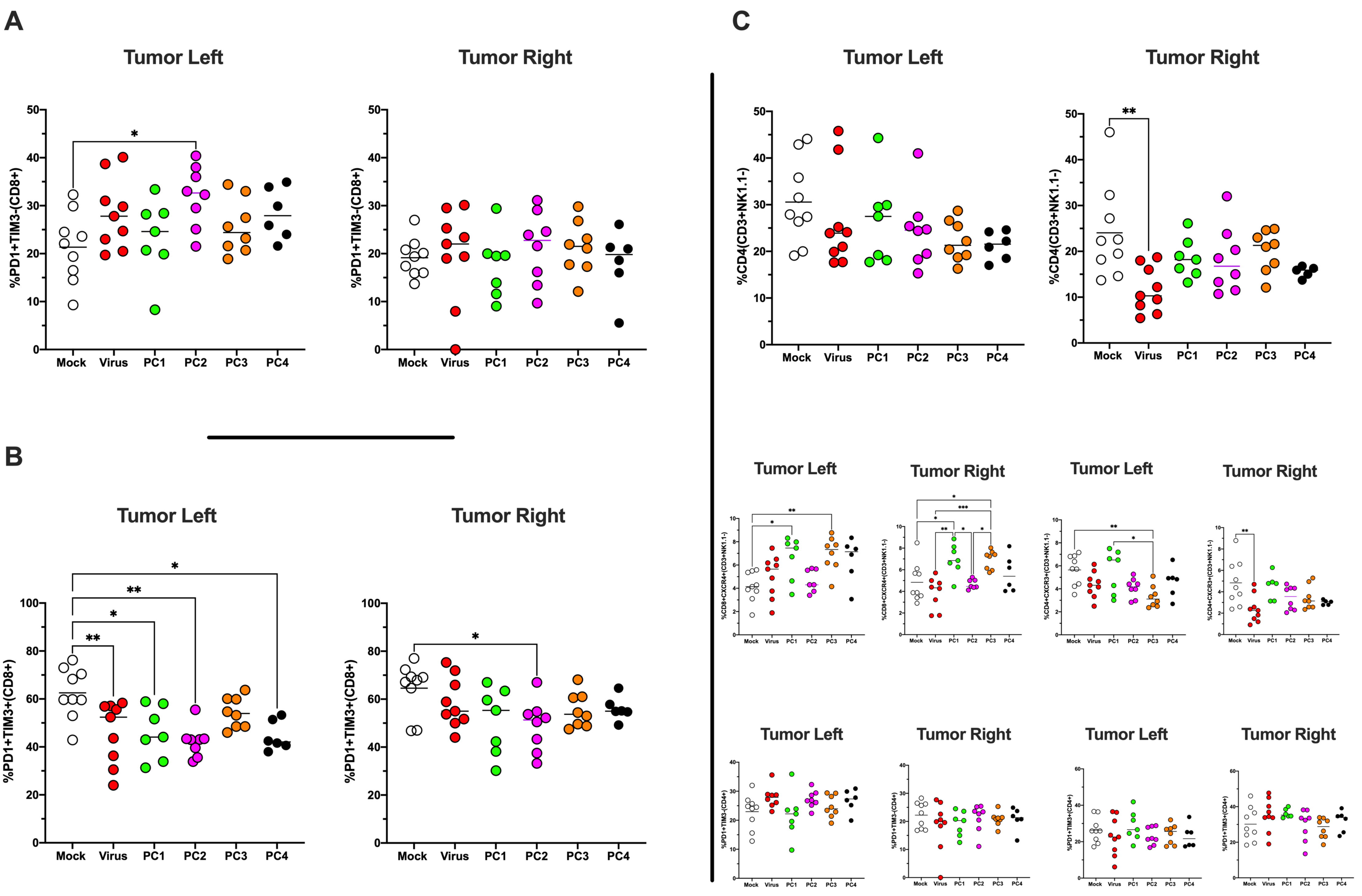
