## Supplementary material for "A novel immunopeptidomic-based pipeline for the generation of personalized oncolytic cancer vaccines": Table 1

| Uniprot ID | Gene name | Peptide sequence | Laumont et al., 2018 |
|---|---|---|---|
| Q64437 | Adh7 | AGASRIIGI | 1 |
| Q9EQH7 | Ndst3 | FYATIIHDL | 0 |
| O08696 | Foxm1 | SGPNRFILI | 1 |
| Q9CXG9 | Phf19 | QGPEYIERL | 1 |
| Q8R3J5 | Chac1 | KYLNVREAV | 0 |
| Q61001 | Lama5 | HYLPDLHHM | 0 |
| Q09143 | Slc7a1 | SYIIGTSSV | 1 |
| O35495 | Cdk14 | SYIHQRYIL | 1 |
| O08784 | Tcof1 | GYMTPGLTV | 0 |
| Q91ZX7 | Lrp1 | SYLIGRQKI | 1 |
| Q61009 | Scarb1 | RGPYVYREF | 0 |
