## Supplementary material for "A novel immunopeptidomic-based pipeline for the generation of personalized oncolytic cancer vaccines": Table 2

| Uniprot ID | Peptide sequence | Pathogen species | Viral Peptide | Laumont et al., 2018 |
|---|---|---|---|---|
| O88738-3 | SYHPALNAI | Molluscum contagiosum virus subtype 1 | SYHAALNAL | 1 |
| Q9QXZ0 | AFHSSRTSL | Human adenovirus A serotype 31 | HFSTSRTSL | 1 |
| Q3TWW8 | SYSDMKRAL | Cercopithecine herpesvirus 1 | AYQDTKRAL | 1 |
| P70452 | NYNSVNTRM | Human herpesvirus 7 | FYNSVNTRN | 0 |
| Q80TP3 | SYLTSASSL | Influenza A virus | TIWTSASSI | 0 |
| Q8VCF0 | SYLPPGTSL | Epstein-Barr virus | TYLPPSTSS | 1 |
| O70405 | FYEKNKTLV | Orf virus | NYYKNKSLV | 0 |
| Q9D1R1 | FYKNGRLAV | Human adenovirus F serotype 41 | AYMNGRVAV | 0 |
| Q91XE7 | KGPNRGVII | Variola virus | KNPNRFVIF | 1 |
| Q6URW6-2 | LYKESLSRL | Human cytomegalovirus | LYLETLSRI | 0 |
| Q9JL70 | RYLPAPTAL | Influenza C virus | RNMPAATAL | 1 |
| O54692 | KYIPAARHL | Human cytomegalovirus | SHQPAARRL | 1 |
| P54775 | YYVRILSTI | Molluscum contagiosum virus subtype 1 | YVFRLLSTI | 1 |
| P54775 | SYRDVIQEL | Human cytomegalovirus | RYADVIQEV | 0 |
| Q61036 | KFYDSKETV | Human adenovirus A serotype 18 | NFYNSKETV | 1 |
