## Supplementary material for "A novel immunopeptidomic-based pipeline for the generation of personalized oncolytic cancer vaccines": Table 3

| Group 1 | Group 4 | Group 7 |
|---|---|---|
| 1. SYHPALNAI | 10. FYEKNKTLV | 18. SGPNRFILI |
| 2. SYLTSASSL | 11. KGPNRGVII | 19. SYIIGTSSV |
| 3. YYVRILSTI | 12. FYKNGRLAV | 20. RGPYVYREF |
| **Group 2** | **Group 5** | **Group 8** |
| 4. SYLPPGTSL | 13. LYKESLSRL | 21. FYATIIHDL |
| 5. RYLPAPTAL | 14. SYRDVIQEL | 22. GYMTPGLTV |
| 6. KYIPAARHL | 15. KFYDSKETV | 23. SYLIGRQKI |
| **Group 3** | **Group 6** | **Group 9** |
| 7. AFHSSRTSL | 16. KYLNVREAV | 24. AGASRIIGI |
| 8. NYNSVNTRM | 17. HYLPDLHHM | 25. QGPEYIERL |
| 9. SYSDMKRAL | | 26. SYIHQRYIL |
