## Supplementary material for "A novel immunopeptidomic-based pipeline for the generation of personalized oncolytic cancer vaccines": Table 4

| Name of peptide | Peptide sequence | Net charge pH 7 | Poly-lysine peptide | Net charge pH 7 |
|---|---|---|---|---|
| Peptide 1 | SYLPPGTSL | 0 | KKKKKKSYLPPGTSL | 6 |
| Ppetide 2 | RYLPAPTAL | 1 | KKKKKKRYLPAPTAL | 7 |
| Peptide 3 | KYIPAARHL | 2.1 | KKKKKKYIPAARHL | 7.1 |
| Peptide 4 | LYKESLSRL | 1 | KKKKKKLYKESLSRL | 7 |
| Peptide 5 | KYLNVREAV | 1 | KKKKKKYLNVREAV | 6 |
| Peptide 6 | FYATIIHDL | -0.9 | KKKKKKKFYATIIHDL | 6.1 |
| Peptide 7 | SPSYAYHQF | 0.1 | KKKKKKSPSYAYHQF | 6.1 |
