## Supplementary material for "A novel immunopeptidomic-based pipeline for the generation of personalized oncolytic cancer vaccines": Table 5

| Peptide | Uniprot ID | Gene name | |
|---|---|---|---|
| KKKKKKSYLPPGTSL | Q8VCF0 | MAVS | PeptiCRAd1 |
| KKKKKKRYLPAPTAL | Q9JL70 | FANCA | |
| KKKKKKYIPAARHL | O54692 | ZW10 | PeptiCRAd2 |
| KKKKKKLYKESLSRL | Q6URW6-2 | MYH14 | |
| KKKKKKYLNVREAV | Q8R3J5 | Chac1 | PeptiCRAd3 |
| KKKKKKKFYATIIHDL | Q9EQH7 | Ndst3 | |
